# Herpes simplex virus 1 subverts the mitochondrial network to support the infection: A lesson on mitochondrial versatility

**DOI:** 10.64898/2026.09.02.748825

**Authors:** Rabina Saud, Kimberly Foster-Lemieur, Brett Duguay, Russell Swerdlow, Maria Kalamvoki

## Abstract

Herpes simplex virus 1 (HSV-1) infects approximately 67% of the population worldwide. It establishes lifelong reservoirs in sensory neurons and has been linked to several diseases including neuronal dysfunction. Disruption of mitochondrial homeostasis is a hallmark of HSV-1 infection, however a molecular understanding of these changes and their significance is not yet well defined. HSV-1 infection causes a UL12.5-dependent inhibition of mitochondrial biogenesis through the loss of mitochondrial DNA and mitochondrial transcription factors, PGC-1α (peroxisome proliferator-activated receptor-gamma co-activator) and TFAM (mitochondrial transcription factor). Conversely, UL12.5-independent mechanisms inhibit mitochondrial fusion by activating the OMA1 metallopeptidase that cleaves the inner mitochondrial membrane fusion protein OPA1 (optic atrophy protein 1) and by down-modulating the outer mitochondrial membrane fusion protein MFN2 (mitofusin 2). This inhibition of fusion results in a smaller mitochondrial network that clusters to perinuclear regions, likely supplying energy for viral replication and envelopment. The inner mitochondrial membrane protein TIM23 is also down-modulated during infection in a UL12.5-independent mechanism. Failure of the virus to promote these changes negatively impacts the infection. Despite these changes, mitochondria are protected from mitophagy due to the viral-induced degradation of several mitophagy adaptor proteins, whereby damaged mitochondrial components, including mitochondrial DNA, are extruded via extracellular vesicles. These mitochondrial changes still support functions necessary for HSV-1 infection. Basal cell respiration is preserved, while spare respiratory capacity and extracellular acidification rates increase, indicating glycolytic activity. Mitochondrial membrane potential is also preserved. Overall, our studies provide mechanistic insight into how HSV-1 impacts mitochondria, which could contribute to viral pathogenesis.

**Importance:** Mitochondria are often referred to as the “powerhouse” of the cell because they are the main energy producers. Disruption of mitochondrial homeostasis is associated with multiple diseases and occurs after infection with pathogens such as HSV-1. By investigating the mechanism(s) by which HSV-1 disrupts mitochondrial homeostasis, we can better understand how HSV-1 causes pathogenesis. HSV-1 infection impacts mitochondrial homeostasis through disruption of four key processes, including: 1) inhibition of mitochondrial biogenesis and the generation of new mitochondria; 2) inhibition of mitochondrial fusion, which rescues reversibly damaged mitochondria; 3) sustaining mitochondrial fission, which removes damaged content; and 4) preventing mitophagy, which clears damaged mitochondria. UL12.5–dependent and UL12.5– independent events during HSV-1 infection disrupt mitochondrial homeostasis, redirecting mitochondrial resources towards progeny virus production. These changes cause irreversible damage to host cells, ultimately driving pathogenesis.

## Introduction

Mitochondria are double-membrane organelles that produce energy in the form of ATP (1–7). They also play significant roles in metabolism, intracellular Ca^2+^ regulation, and in the control of apoptosis (8). Mitochondria have an outer (OMM) and an inner mitochondrial membrane (IMM), which define two aqueous compartments: the intermembrane space (IMS), and the mitochondrial matrix (9). The IMM contains infoldings called cristae. Within the matrix is the mitochondrial genome (10). Mitochondrial homeostasis is maintained through a balance of mitochondrial biogenesis and mitophagy (3;7). Mitochondrial biogenesis, the generation of new mitochondria, is orchestrated by PGC-1α (peroxisome proliferator-activated receptor-gamma co-activator). PGC-1α activates a network of transcription factors, including TFAM (mitochondrial transcription factor), which together regulate mitochondrial DNA (mtDNA) transcription, replication, stability, and packaging in nucleoids (11–14). Mitophagy, involves the removal of damaged mitochondria (15). Mitochondria also continuously undergo fission and fusion, changing their structure, number, and size in order to maintain network integrity (1–7). Mitochondrial fusion allows reversibly damaged mitochondria to exchange gene products and metabolites to restore respiratory capacity. Fusion of the OMM is facilitated by the membrane GTPases, mitofusin 1 and mitofusin 2 (MFN1 and MFN2), whereas fusion of the IMM is enabled by OPA1 (optic atrophy protein 1) (16–20). Conversely, mitochondrial fission occurs when mitochondria divide or are irreversibly damaged; a process that is coordinated by dynamin-related protein 1 (Drp1), following its recruitment to the OMM by mitochondrial fission protein 1 (Fis1) (21;22). Mitochondria are the “powerhouse” of the cell. Enzymatic reactions in the matrix, known as tricarboxylic acid (TCA) cycle, transfer electrons through the electron transport chain (ETC) to create the electrochemical gradient that drives ATP generation through oxidative phosphorylation (OXPHOS) (3;7;23).

Previous studies have determined that HSV-1 infection disrupts mitochondrial homeostasis by depleting mtDNA and mtRNA. This process is mediated by UL12.5, an N-terminus truncated form of the HSV-1 alkaline deoxyribonuclease UL12, however, its weak nuclease activity is not involved (24;25);(26). Instead, it was proposed that the virus may employ host mitochondrial nucleases, such as endonuclease G (ENDOG) and endonuclease G-like 1 (EXOG), but depleting these nucleases only partially restores mtDNA levels (26). Recently, it was reported that inhibition of mtDNA replication results in enlarged mitochondrial nucleoids that are typically eliminated through an endolysosomal pathway (27). This also occurs following UL12.5 expression, but the extent to which this mechanism is responsible for mtDNA down-modulation is unclear (27).

Here, we sought to delineate the events that disrupt mitochondrial homeostasis in HSV-1–infected cells. We observed that the reduction in mtDNA levels during infection coincides with decreased levels of TFAM and PGC-1α. Typically, changes in TFAM protein levels cause proportional alterations in mtDNA levels, resulting in impaired mitochondrial biogenesis. We also found that HSV-1 infection disrupts mitochondrial fusion by down-modulating MFN2 and activating the mitochondrial metalloendopeptidase OMA1. After activation, OMA1 cleaves OPA1 into soluble, non-membrane-associated forms that do not support fusion (20;28). Mitochondrial fission, on the other hand, appears to be sustained. One hypothesis for why HSV-1 infection disables fusion but sustains fission is for promoting the formation of smaller mitochondria that are easier to translocate to perinuclear sites where they likely provide energy for virus replication and virion envelopment (29–33). In support of this hypothesis, we found that blocking the ability of the virus to inhibit mitochondrial fusion by depleting OMA1 or by expressing a long, membrane-associated form of OPA1 resulted in delayed viral gene expression and decreased progeny virus production. We observed a similar delay in viral gene expression when we inhibited mitochondrial fission. In addition, translocase of inner mitochondrial membrane 23 (TIM23), a key component of the homonymous IMM translocase complex, is depleted in HSV-1-infected cells (34;35). This is important during infection, as failure of the virus to deplete TIM23 caused a delay in viral gene expression. Despite these changes, we observed an increased spare respiratory capacity during HSV-1 infection, and a dependence of the infection on glycolysis (36–38). Also, we found that the mitochondrial membrane potential is maintained until the late stages of infection, and ROS production is not stimulated. Altogether, these findings support a strategic viral mechanism whereby mitochondria are altered to support the infection. To further support this model, we found that major mitophagy adaptors are lost during the early stages of infection, likely protecting the altered mitochondria from elimination (39). We also documented the selected removal of mitochondrial factors during infection, including mtDNA and mtDNA-binding proteins, via extracellular vesicles (EVs). Both UL12.5-dependent and UL12.5–independent mechanisms underline mitochondrial homeostasis disruption during HSV-1 infection. Degradation of mtDNA and TFAM is UL12.5-dependent (24;25), (26). Conversely, conversion of OPA1 to soluble forms, TIM23 down-modulation, and the loss of mitophagy adaptors are independent of UL12.5. Finally, we determined that UL12.5 expression, despite disrupting mitochondrial homeostasis, did not activate innate immune responses in human cells where the virus productively replicates, rather it sensitized the cells, priming their immune-sensing pathways.

These findings demonstrate that disruption of mitochondrial homeostasis during HSV-1 infection arises from multiple convergent events. Here we provide key mechanistic insight into how mitochondria are modified in HSV-1-infected cells, highlighting the metabolic interplay during infection, which overall advances our understanding of the molecular underpinning in virus-driven pathogenesis.

## Results

### HSV-1 infection inhibits mitochondrial fusion but preserves fission

Mitochondrial fusion is a two-step process; the first step requires the fusion of the OMM and is mediated by MFN1 and MFN2, which have non-redundant roles (40;41). The second step requires fusion of the IMM and is mediated by OPA1 (40;41). The expression of MFN1/2 and OPA1 was assessed during HSV-1 infection. We infected hTERT-HEL cells with either HSV-1(F) or a ΔUL47 virus (2.5 PFU/cell). The ΔUL47 virus has a delayed onset of beta- and gamma-gene expression due to: 1) The absence of the UL47 transactivation function; 2) Defects in the viral kinase Us3, which has anti-apoptotic roles; and 3) Defects in the viral RNase VHS (virion host shutoff), which degrades host mRNAs for immune evasion (42) (43;44). Additionally, ΔUL47 virus displays growth defects, since it activates type I IFN and pro-inflammatory responses and cannot evade autophagy (Figure S1). As a result, we expect extensive mitochondrial damage in ΔUL47 virus-infected cells, whereas wild type (WT) virus is expected to preserve certain mitochondria functions. When we analyzed the levels of MFN1 and MFN2 over time, we found that MFN1 remained unaltered during WT-virus infection but decreased in ΔUL47 virus-infected cells, with the concomitant appearance of a higher migrating form of unknown origin (Figure 1A). This band could represent a modified MFN1 form, which appears when mitochondrial fusion status changes (45;46). MFN2, on the other hand, was decreased by both viruses at late times post-infection (Figure 1A and quantification).

**Figure 1:**
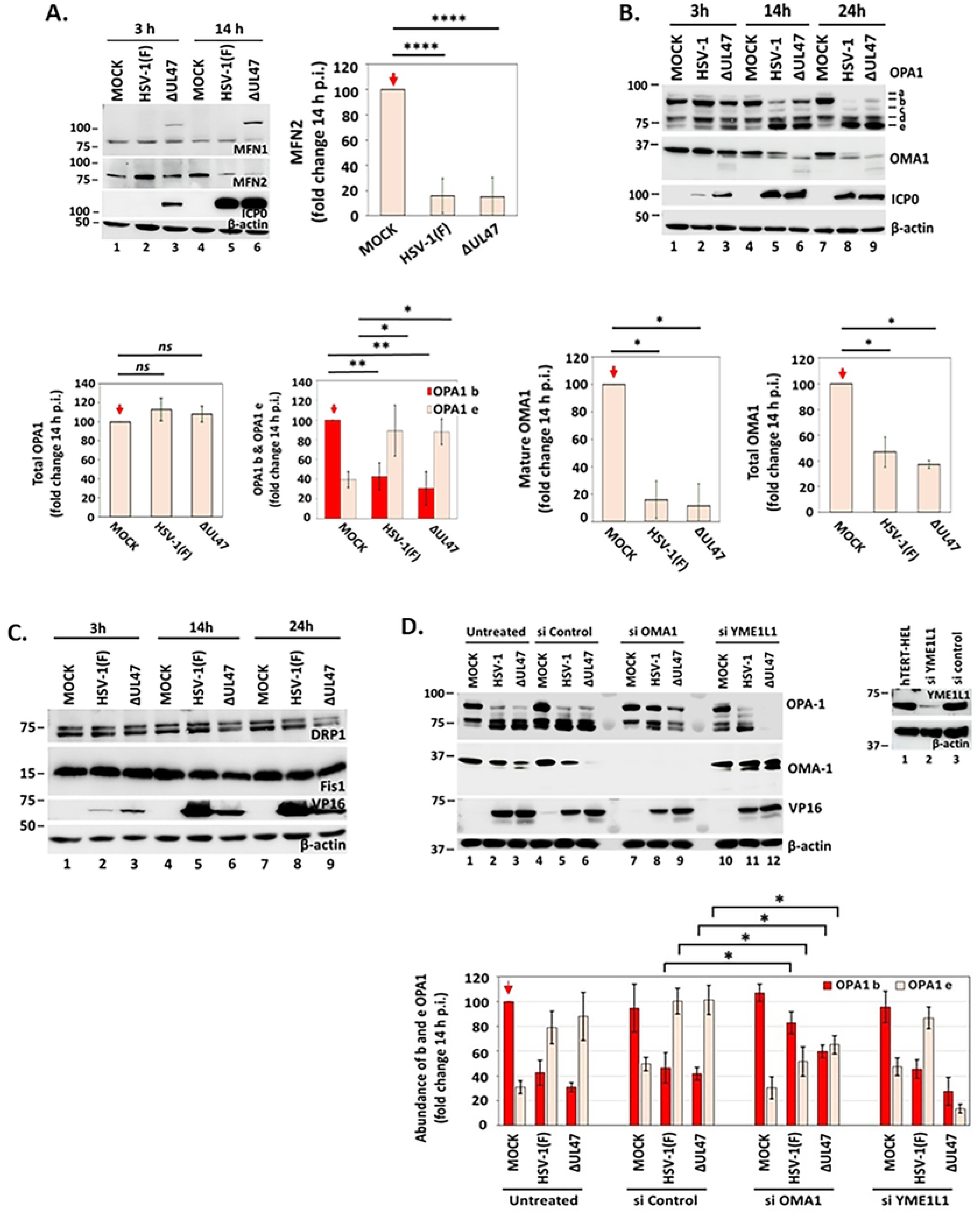
HSV-1 infection modifies OMM and IMM proteins. **A.** hTERT-HEL cells were infected with HSV-1(F) and ΔUL47 virus (2.5 PFU/cell). The cells were harvested at 3 and 14 hpi and equal amounts of proteins from total cell lysates were analyzed for MFN1 and MFN2. ICP0 served as a control for infection and β-actin as a loading control. Quantification of MFN2 protein signal from three independent experiments was done using ImageJ. P<0.0001**** with two-tailed Student’s *t*-test. **B.** hTERT-HEL were infected with HSV-1(F) and ΔUL47 virus (2.5 PFU/cell). The cells were harvested at 3, 14, and 24 hpi and equal amounts of proteins from total cell lysates were analyzed for OPA1 and OMA-1. ICP0 served as a control for infection and β-actin as a loading control. Quantifications of total OPA1, b and e forms of OPA1, mature OMA1, and total OMA1 from three independent experiments was done using ImageJ. P<0.05*, P<0.01** with two-tailed Student’s *t*-test. **C.** hTERT-HEL cells were infected with HSV-1(F) and ΔUL47 virus (2.5 PFU/cell). The cells were harvested at 3, 14, and 24 hpi and equal amounts of proteins from total cell lysates were analyzed for DRP1 and Fis1. VP16 served as a control for infection and β-actin as a loading control. **D.** hTERT-HEL cells were transfected with an OMA1 siRNA, YME1L1 siRNA, scrambled siRNA (200 ng/10^6^ cells), or left untransfected. At 72 h post-transfection the cells were infected with HSV-1(F) and ΔUL47 virus (10 PFU/cell). The cells were harvested at 16 hpi and equal amounts of proteins from total cell lysates were analyzed for OPA1, OMA1, and YME1L1. VP16 served as a control for infection and β-actin as a loading control. Signal quantification for total OPA1, as well as b and e forms of OPA1 from three independent experiments was done using ImageJ. Statistical significance was evaluated with two-tailed Student’s *t*-test. P<0.05*, P<0.01**, P<0.001***, P<0.0001****, ns= not significant.

Human *OPA1* is alternatively spliced to generate eight isoforms, collectively known as long (L)-OPA1, which are membrane-bound (16–20). All OPA1 isoforms can be cleaved to generate short (S)-OPA1 forms, which localize into the IMS and are not bound to membranes. L-OPA1 is essential for fusion, mediated by its interaction with MFN1/2, whereas S-OPA1 generates a scaffolding structure, which wraps cristae, and is necessary for anchoring the mtDNA and organizing the respirasome. S-OPA1 is sufficient to maintain cristae morphology and is more efficient than L-OPA1 at maintaining energetic competence, but it cannot support fusion (16–20). In cells infected with WT or ΔUL47 virus, we observed a gradual decrease of L-OPA1 (particularly form b) and an enrichment of S-OPA1 (particularly form e), whereas total OPA1 remained unaltered (Figure 1B and quantifications). These results were reproducible in human primary epidermal keratinocytes and HaCaT keratinocytes (Figure 5B and S3A). The levels of proteins involved in fission (Drp1 and Fis1) remained unaltered in WT virus infections, while Drp1 levels decreased with ΔUL47 virus (Figure 1C). These findings indicate that the proteins involved in mitochondrial fusion are disrupted in WT virus infected cells and may disproportionally affect the ability of mitochondria to perform mitochondrial fusion.

Two IMM proteases, YME1L1 (YME1-like protein 1) and OMA1, cleave OPA1 to regulate mitochondrial fusion (20;28). OMA1 shows little activity under physiological conditions, but under stress is activated to cleave L-OPA1. YME1L1 adjusts the L-OPA1 to S-OPA1 ratio under physiological conditions, which is required for fusion (20;28). To determine whether OMA1 processes L-OPA1 during HSV-1 infection, we performed two sets of experiments: 1) We analyzed OMA1 levels during HSV-1 infection. OMA1 activation is associated with autocatalytic degradation, which initiates from both ends of OMA1 and results in complete turnover (47). As shown in Figures 1B (and quantification), the levels of mature OMA1 as well as the total OMA1 (after proteolytic processing) were reduced in infected cells, indicating OMA1 activation; and 2) We transiently depleted cells of either OMA1 or YME1L1 and assessed L- to S-OPA1 conversion during HSV-1 infection. As shown in Figure 1D (and quantification), depletion of OMA1 suppressed the conversion of L- to S-OPA1. Conversely, depletion of YME1L1 enhanced the conversion of L- to S-OPA1, likely because YME1L1 depletion stabilizes OMA1 (Figure 1D). In ΔUL47 virus infections where autophagy likely clears damaged mitochondria, OPA1 levels were substantially lower compared to WT virus infections, under both conditions. The efficiency of OMA1 and YME1L1 depletion is shown in Figures 1D. Overall, these data indicate that OMA1 is activated in infected cells, converting L- to S-OPA1, and that YME1L1 negatively regulates OMA1 to prevent OPA1 loss.

To demonstrate that mitochondrial fusion is inhibited in HSV-1 infected cells, we analyzed the mitochondrial network in infected and uninfected cells (5 PFU/cell, 16 h post-infection) using ImageJ and the MiNA plugin (Figure 2A) (48). We found that the mitochondrial footprint (total area occupied by mitochondria) and the mean of the summed branch lengths (the total length of branches in μm in an individual network) were significantly decreased in HSV-1(F)-infected cells, whereas the number of network branches only slightly decreased. Mitochondrial mass also decreased in HSV-1(F)–infected cells (Figure 2A). These observations are consistent with the inhibition of mitochondrial fusion, indicating that mitochondria in infected cells are smaller.

**Figure 2:**
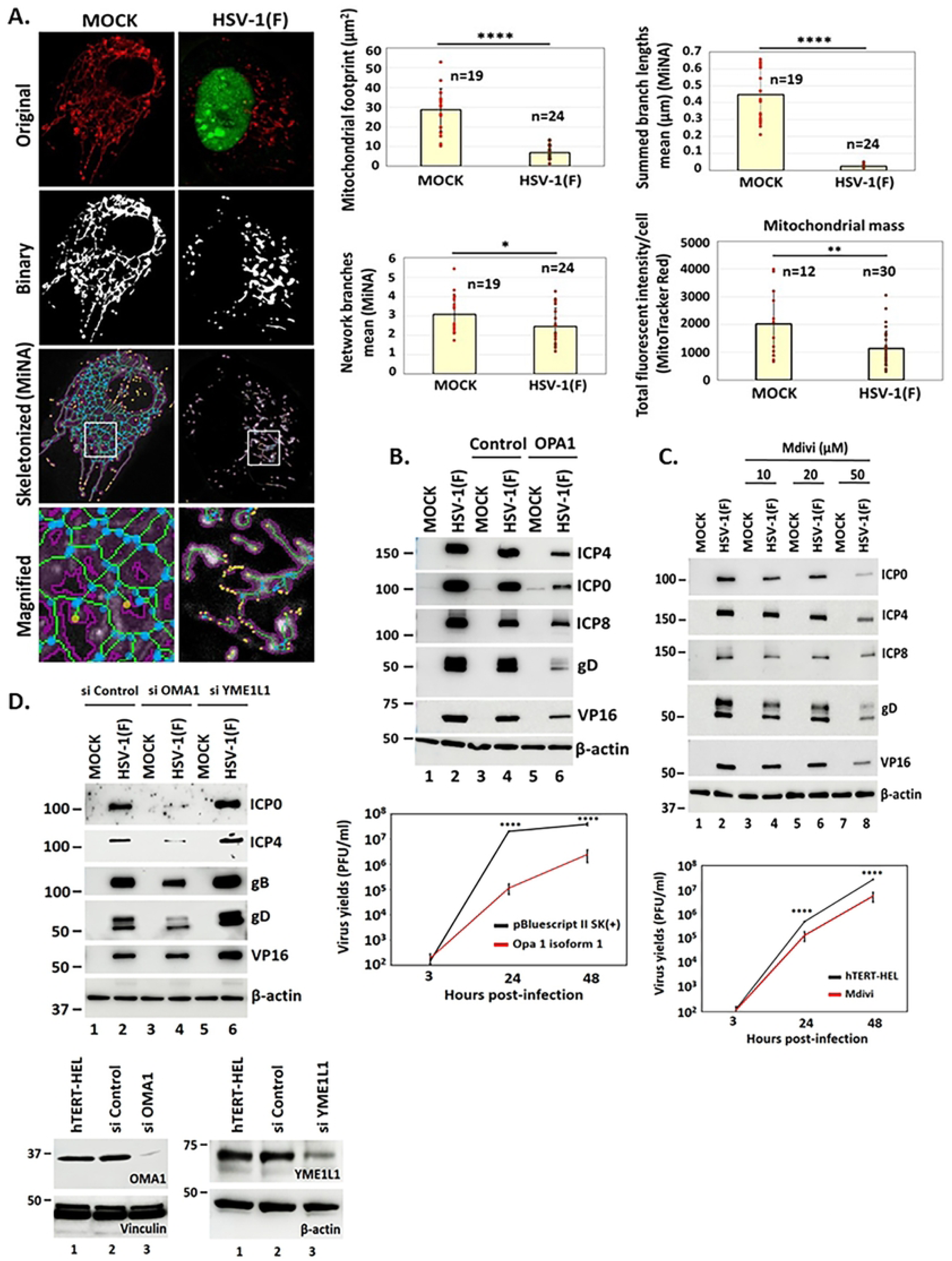
Sustained mitochondrial fusion or inhibition of mitochondrial fission inhibit HSV-1 infection. **A.** hTERT-HEL cells were infected with HSV-1(F) (5 PFU/cell) or left uninfected. Samples were stained with MitoTracker Red (100 nM), fixed at 16 hpi, and immunostained with an ICP4 antibody to visualize the infected cells. Images were acquired with the same settings of the Leica TCS SP8 confocal microscope and analyzed with FiJi and the MiNA plugin for mitochondrial footprint, network branches, summed branch lengths, and with ImageJ for mitochondrial mass (48). Statistical significance was calculated using two-tailed Student’s *t*-test. P<0.05*, P<0.01**, P<0.001***. **B.** ARPE-19 cells seeded in 6 x well plates were transfected with an OPA1-expressing plasmid or control pBluescript II SK(+) (1 μg). At 18 h post-transfection the cells were infected with HSV-1(F) (1 PFU/cell) or left uninfected. The cells were harvested at 12 hpi and equal amounts of proteins from total cell lysates were analyzed for ICP4, ICP0, ICP8, gD, VP16, and β-actin that served as a loading control. In addition, replicate cultures of ARPE-19 cells transfected with the OPA1-expressing plasmid and the control were infected with HSV-1(F) (0.01 PFU/cell). The cells were harvested at 3, 24, and 48 h post-infection and progeny virus production was determined by plaque assays in Vero cells. **C.** hTERT-HEL cells were treated with different doses of Mdivi (10, 20, 50 μM) for 2 h prior to infection with HSV-1(F) (0.5 PFU/cell). The cells were harvested at 12 hpi and equal amounts of proteins from total cell lysates were analyzed as above. In addition, we analyzed HSV-1 growth in the presence of Mdivi (20 μM), and progeny virus production was quantified at 3, 24, and 48 h post-infection. **D.** hTERT-HEL cells were transfected with an OMA1 siRNA, YME1L1 siRNA, scrambled siRNA (200 ng/10^6^ cells), or left untransfected. At 72 h post-transfection the cells were infected with HSV-1(F) (1 PFU/cell). The cells were harvested at 24 hpi and equal amounts of proteins from total cell lysates were analyzed for ICP0, gD, VP16, and β-actin. Efficiency of OMA1 and YME1L1 depletion at 72 h post-transfection is depicted.

Finally, we asked how changes in mitochondrial fusion and fission impact the infection. To assess functional importance, we expressed the isoform 1 of OPA1 in ARPE-19 cells, a long, membrane-anchored OPA1 isoform, and analyzed its impact on HSV-1 infection. This long OPA1 is expected to antagonize the virus-mediated inhibition of mitochondrial fusion. We found that overexpression of OPA1 isoform 1 delays HSV-1 gene expression and causes a decrease in progeny virus production by at least 100-fold (Figure 2B). We also treated hTERT-HEL cells with Drp1 inhibitor to suppress fission and observed a dose-dependent inhibition of viral gene expression and at least 5-fold decrease in virus yields with Mdivi at 20 μM (Figure 2C). Finally, we depleted cells of either OMA1, which is required for OPA1 conversion, or YME1L1, which suppresses OMA1 activity. We observed that the depletion of OMA1 causes a delay in viral gene expression, whereas the loss of YME1L1 enhances viral gene expression (Figure 2D). Together, these data indicate that HSV-1 infection inhibits mitochondrial fusion, which is functionally significant as stabilization of mitochondrial fusion or inhibition of mitochondrial fission negatively impact the infection.

### HSV-1 infection down-modulates mitochondrial biogenesis factors

Mitochondrial fusion is crucial for mitochondrial biogenesis and for integrating new mitochondria into the network (40;49). Since S-OPA1 interacts directly with mtDNA, impacting TFAM localization, we wanted to understand the role of TFAM during infection (16;18;50;51). TFAM is a mitochondrial transcription factor that plays an essential role in mtDNA transcription, replication, and packaging in nucleoids. The protein levels of TFAM were monitored during HSV-1 infection of hTERT-HEL. As shown in Figure 3A (and the quantification) a reduction of TFAM was observed as soon as 8 h post-infection that became apparent at 10 h post-infection, which is consistent with previous published data in mouse cells (52). Additionally, the conversion of L- to S-OPA1 was observed as soon as 6 h post-infection (Figure 3A). Since TFAM is down-modulated in HSV-1–infected cells, we next asked how TFAM overexpression would impact HSV-1 infection. We developed an hTERT-HEL cell line, stably expressing exogenous TFAM (TFAM-HEL), by transducing the cells with a lentiviral vector carrying the TFAM open reading frame (ORF) under CMV promoter, followed by puromycin selection. TFAM-HEL and the parental cell line (hTERT-HEL) were infected with HSV-1(F) (0.01 PFU/cell), and progeny virus production was quantified by plaque assays at 3, 24, and 48 h post-infection. We found no differences in virus yields between the two cell lines (Figure 3B). This is likely because HSV-1 infection efficiently down-modulates both exogenous and endogenous TFAM (Figure 3C, 3D). TFAM reduction likely occurs at the protein level, as *TFAM* mRNA levels between infected and uninfected cells were similar (Figure S2A). Since TFAM levels directly correlate with mtDNA levels, we also quantified mtDNA in TFAM-HEL and hTERT-HEL cells infected with the WT virus at 14 h. As shown in Figure 3E, mtDNA levels were decreased in both cell lines with infection. The different molecular weights of TFAM in the TFAM-HEL likely represent the non-cleaved and a cleaved (mature) TFAM form after removal of a 43 aa mitochondrial localization sequence (MLS) at the N-terminus of the protein.

**Figure 3:**
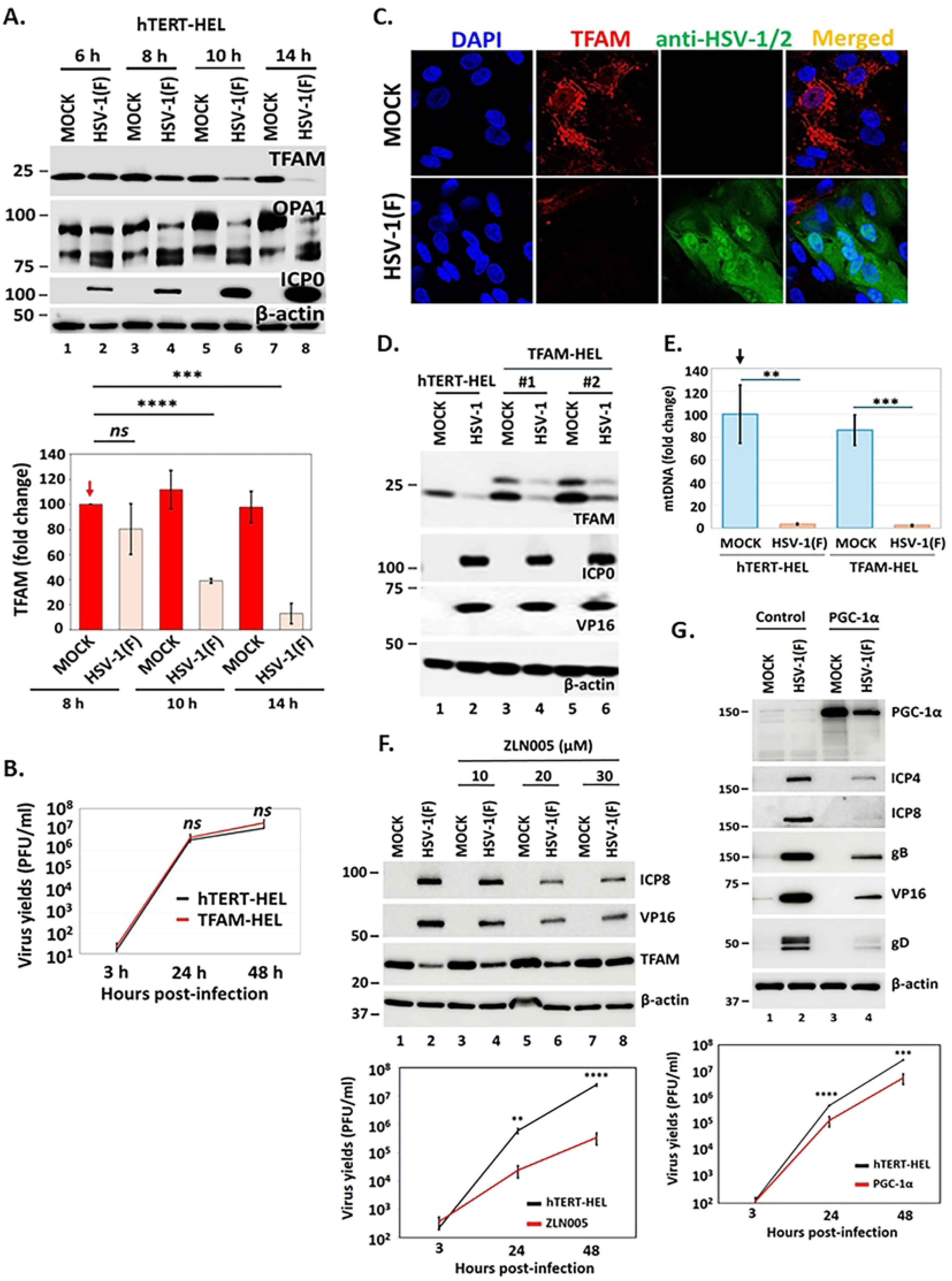
HSV-1 infection down-modulates mitochondrial transcription factors. **A.** hTERT-HEL cells were infected with HSV-1(F) (5 PFU/cell). The cells were harvested at 6, 8, 10, and 14 hpi and equal amounts of proteins from total cell lysates were analyzed for TFAM and OPA1. ICP0 served as a control for infection and β-actin as a loading control. Quantification of TFAM protein signal from three independent experiments is depicted. P<0.001***, P<0.0001**** with Student’s *t*-test. **B.** hTERT-HEL and TFAM–expressing hTERT-HEL cells (TFAM-HEL) were infected with HSV-1(F) (0.01 PFU/cell). The cells were harvested at 3, 24, and 48 hpi and progeny virus production was quantified by plaque assays. Three biological replicates were used. **C.** Replicate cultures of the TFAM-HEL cell line were infected with HSV-1(F) (2 PFU/cell) or left uninfected. The cells were fixed at 12 hpi and stained with TFAM and an HSV-1/2 antibody. Images were acquired with the same settings of a Leica TCS SP8 confocal microscope. **D.** hTERT-HEL cells and two different clones of TFAM-expressing derivatives were infected with HSV-1(F) (2 PFU/cell) or left uninfected. The cells were harvested at 14 hpi and equal amounts of proteins from total cell lysates were analyzed for TFAM. ICP0 and VP16 served as controls for infection and β-actin as a loading control. **E.** Infections were same as in panel D. Analysis of mtDNA was done at 14 hpi by qPCR analysis. Statistical significance was determined by two-tailed Student’s *t*-test using three biological replicates. P<0.01**, P<0.001***. **F.** hTERT-HEL cells were treated for 2 h with different doses of ZLN005 or left untreated. The cells were then infected with HSV-1(F) (2.5 PFU/cell), and protein analysis was performed at 12 hpi using equal amounts of proteins from total cell lysates. Growth curves were performed using 10 μM of ZLN005 and infection with HSV-1(F) at 0.01 PFU/cell. **G.** ARPE-19 cells were transfected with a PGC-1a-expressing plasmid or the control pBluescript II SK(+). At 24 h post-transfection the cells were infected with HSV-1(F) (0.5 PFU/cell). Samples were harvested at 12 hpi and equal amounts of proteins from total cell lysates were analyzed for viral proteins. Growth curves were performed under the same conditions.

We also monitored transcription factors and regulators upstream of TFAM to determine if they were impacted during HSV-1 infection. We found that PGC-1α and its splicing variants were down-modulated in infected cells, likely at the protein level, as mRNA levels were comparable between infected and uninfected cells (Figure S2A, S2B and quantification). To determine why HSV-1 down-modulates PGC-1α, we treated cells with ZLN005 that functions as a transcriptional activator of PGC-1α, followed by HSV-1 infection. We observed a dose-dependent inhibition of HSV-1 gene expression (Figure 3F, S2C), along with delayed TFAM degradation (Figure 3F). Also, progeny virus production was decreased by 100-fold in the presence of ZLN005 (10 μM). In a complementary approach, we expressed PGC-1α exogenously, followed by HSV-1 infection. We observed that exogenous PGC-1α delayed viral gene expression and progeny virus production (Figure 3G), albeit to a lower extent compared to ZLN005. This difference could be because exogenous expression of PGC-1α does not necessarily means activation. It could also mean that ZLN005 has off-target effects. These data suggest a mechanism whereby the virus down-modulates the mitochondrial transcription factor PGC-1α, likely to evade its negative impact on infection.

Finally, we analyzed several other mitochondrial transcription factors, both downstream and upstream of PGC-1α. We found that the nuclear respiratory factor 1 (NRF1), which is downstream of PGC-1α, was not altered between WT-virus–infected and uninfected cells. There was, however, a reduction in ΔUL47 -infected cells, with the concomitant appearance of a shorter product of unknown origin (Figure S2B). The levels of sirtuin 1 (SIRT1), a deacetylase which regulates PGC-1α, were also unaltered between infected and uninfected cells (Figure S2B) (53). This is likely because SIRT1 is required for optimal virus yields, as determined after comparing HSV-1 growth in hTERT HEL versus SIRT1 KD derivatives (Figure S2D). Finally, the Yin Yang (YY1) transcription factor that is upstream of SIRT1 was marginally upregulated in HSV-1 –infected cells (Figure S2E). This factor is also required for optimal virus yields, as demonstrated in YY1 KD cells, where viral gene expression and progeny virus production were reduced (Figure S2F).

Together these data indicate that key mitochondrial biogenesis factors such as PGC-1α and TFAM are down-modulated in HSV-1-infected cells, likely because they negatively impact the infection, whereas upstream transcriptional factors are harnessed by the virus to support viral gene expression.

### HSV-1 selectively down-modulates components of IMM translocase complexes

Down-modulation of MFN2 and conversion of L-OPA1 to S-OPA1 impact the OMM and IMM, respectively. Thus, we investigated how the TOM (translocase of the outer mitochondrial membrane) and TIM (translocase of the inner mitochondrial membrane) multi-subunit complexes that import proteins from the cytoplasm into the mitochondria are impacted by HSV-1 infection (54;55). We performed a kinetic analysis to monitor the levels of different components of these complexes, including TOM20, TOM40, TOM70, and TIM23. We found that the levels of most TOM proteins remained unaltered between WT-virus– infected and uninfected cells (Figure 4A). Some potential degradation of TOM40 was observed exclusively in ΔUL47 virus–infected cells, as soon as 3 h post-infection, appearing as a smear (Figure 4A). In contrast, TIM23 was decreased by both viruses (Figure 4A and quantification).

**Figure 4:**
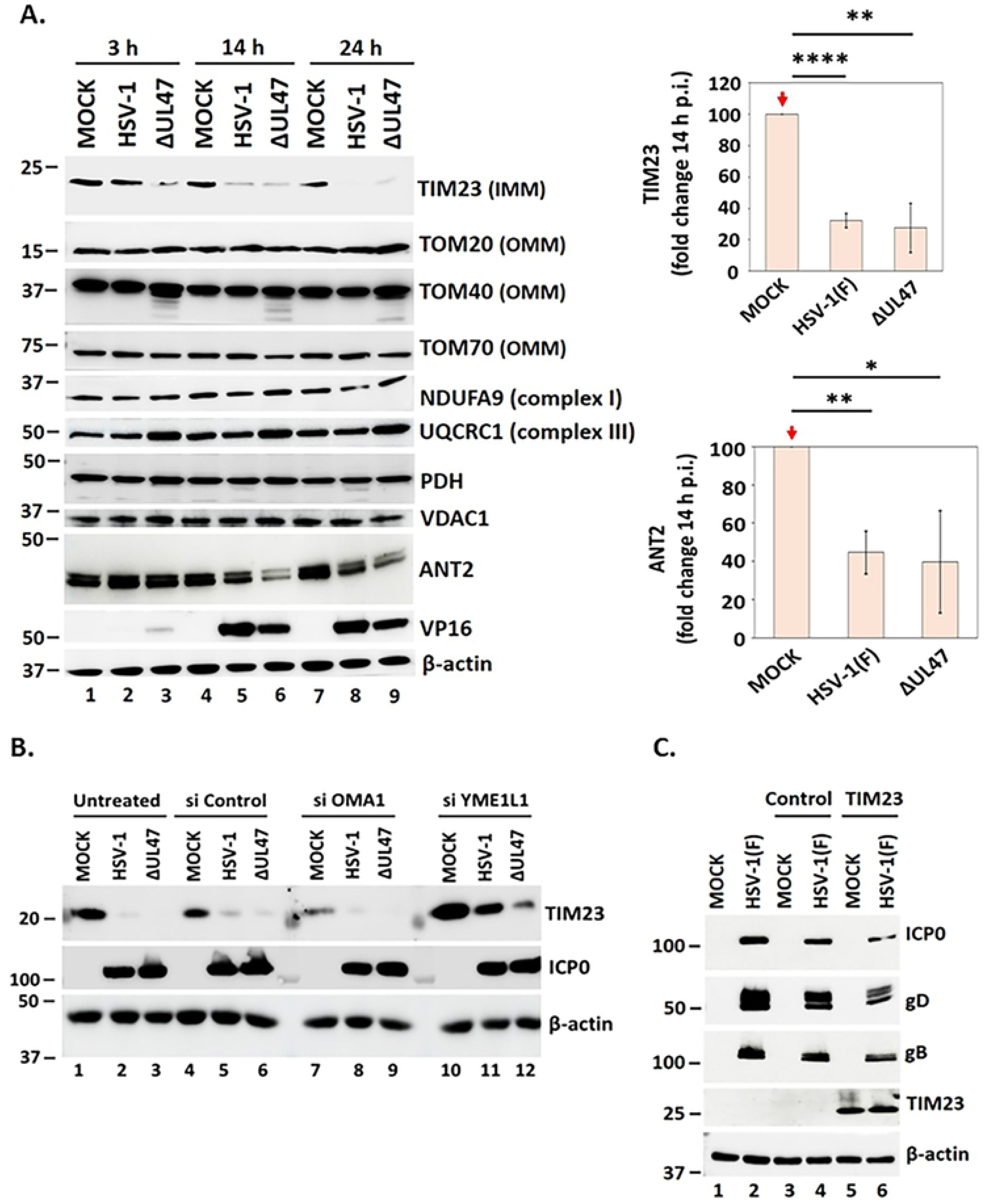
IMM proteins are down-modulated in HSV-1 infected cells. **A.** hTERT-HEL cells were infected with HSV-1(F) and ΔUL47 virus (2.5 PFU/cell) or left uninfected. The cells were harvested at 3, 14, and 24 hpi and equal amounts of proteins from total cell lysates were analyzed for TIM23, TOM20, TOM40, TOM70, NDUFA9, UQCRC1, PDH, VDAC1, and ANT2. VP16 served as a control for infection and β-actin as a loading control. Quantification of TIM23 and ANT2 protein signals from three independent experiments was done using ImageJ. Statistical significance was determined by two-tailed Student’s *t*-test using three biological replicates. P<0.05*, P<0.01**, P<0.0001***, ns=not significant. **B.** hTERT-HEL cells were transfected with an OMA1 siRNA, YME1L1 siRNA, scrambled siRNA (200 ng/10^6^ cells), or left untransfected, as in Figure 1D. At 72 h post-transfection the cells were infected with HSV-1(F) and ΔUL47 virus (10 PFU/cell). The cells were harvested at 16 hpi and equal amounts of proteins from total cell lysates were analyzed for TIM23. ICP0 served as a control for infection and β-actin as a loading control. Efficiency of OMA and YME1L1 depletion is as in Figure 1D. **C.** ARPE-19 cells were transfected with a TIM23-expressing plasmid or control pBluescript II SK(+) for 18 h followed by infection with HSV-1(F) (1 PFU/cell). Viral gene expression was analyzed at 16 hpi.

We also analyzed the levels of VDAC1 (voltage-dependent anion channel 1) and ANT2 (adenine nucleotide translocase 2) that function in the outer and inner mitochondrial membrane, respectively (56–58). Evidence supports a functional interaction between VDAC and ANT isoforms to facilitate metabolite channeling between the cytoplasm and mitochondrial matrix (59). We did not detect any change in the levels of VDAC1 between infected and uninfected cells, but we did observe a decrease in ANT2 protein levels at late times post-infection (Figure 4A and quantification). During oxidative phosphorylation, ANT2 mediates the exchange of ATP, synthesized in the mitochondrial matrix, with ADP from the cytoplasm (57). During glycolysis, ANT2 imports ATP to mitochondria from the cytoplasm (57). ANT2 also functions as a mitochondrial RNA transport translocon (60). This ANT2-mediated efflux of mt-dsRNA is the primary trigger of immune responses (60). It is therefore possible that HSV-1 infection down-modulates ANT2 to evade these antiviral responses, and to prevent ATP uptake from the cytoplasm that will be used during infection.

Further, we monitored the levels of different electron transport chain (ETC) components including UQCRC1 (Ubiquinol-Cytochrome C Reductase Core Protein 1), NDUFA9 (NADH:Ubiquinone Oxidoreductase Subunit A9), and PDH (pyruvate dehydrogenase). We observed that these proteins remained unaltered in HSV-1-infected cells (Figure 4A). These proteins may be stable due to their long half-life, and this is critical for the functionality of mitochondria during infection (25;32);(61).

We also observed that TIM23 protein levels are regulated by the OMA1 and YME1L1 proteases. OMA1 depletion caused TIM23 down-modulation, whereas depletion of YME1L1 upregulated TIM23 (Figure 4B). This is likely because YME1L1 negatively regulates the TIM23 complex by degrading certain subunits (62). In both cases HSV-1 infection further down-modulated TIM23 (Figure 4B). Given the antagonistic roles of OMA1 and YME1L1 in mitochondrial dynamics, along with the regulation of both proteases by HSV-1, it is likely that these proteins also play an additional role in regulating TIM23 levels during infection. Finally, we exogenously expressed TIM23 in ARPE-19 cells and determined that it has a negative effect on HSV-1 infection (Figure 4C), indicating that TIM23 down-modulation causes mitochondrial changes favorable for the infection.

Overall, HSV-1 infection down-modulates selected mitochondrial transport complexes but retains the levels of several respiratory chain factors to support the infection.

### HSV-1 UL12.5 is required for mtDNA and TFAM degradation, but not for L-OPA1 to S-OPA1 conversion or TIM23 down-modulation

The HSV-1 *UL12* gene encodes two separately promoted 3’ co-terminal mRNAs, which produce two distinct but related proteins: UL12, which localizes to the nucleus, and UL12.5, which localizes to mitochondria (25;63-65). ANF-1 is an HSV-1 mutant that expresses UL12.5, but not UL12, and degrades mtDNA and TFAM, like the WT virus (KOS37 or HSV-1(F)) (Figure 5A-C) (25;63-65). This was not the case with AN-1, a UL12/UL12.5 null virus, or KOS UL98 a virus that expresses the HCMV ortholog of UL12 and does not degrade mtDNA (Figure 5A-C) (25;63-66). These results were expected, as TFAM levels directly correlate with mtDNA levels (67);(52). However, the conversion of L-OPA1 to S-OPA1, as well as TIM23 degradation, occurred independent of TFAM degradation and required neither UL12.5 nor UL12 (Figure 5A-C). These findings suggest that disruption of mitochondrial homeostasis in HSV-1–infected cells result from a series of convergent events, of which the degradation of mtDNA and TFAM are UL12.5-dependent. These findings are reproducible across different cell types where the virus productively replicates, including human immortalized fibroblasts (Figure 5A), human primary epidermal keratinocytes (Figure 5B), human immortalized keratinocytes (Figure S3A), and human immortalized retinal epithelial cells (data not shown). Notably, inhibition of LonP1, a protease that is known to degrade TFAM, resulted in some TFAM accumulation particularly in higher doses of the inhibitor that also caused a delay in viral gene expression (Figure S3B).

**Figure 5:**
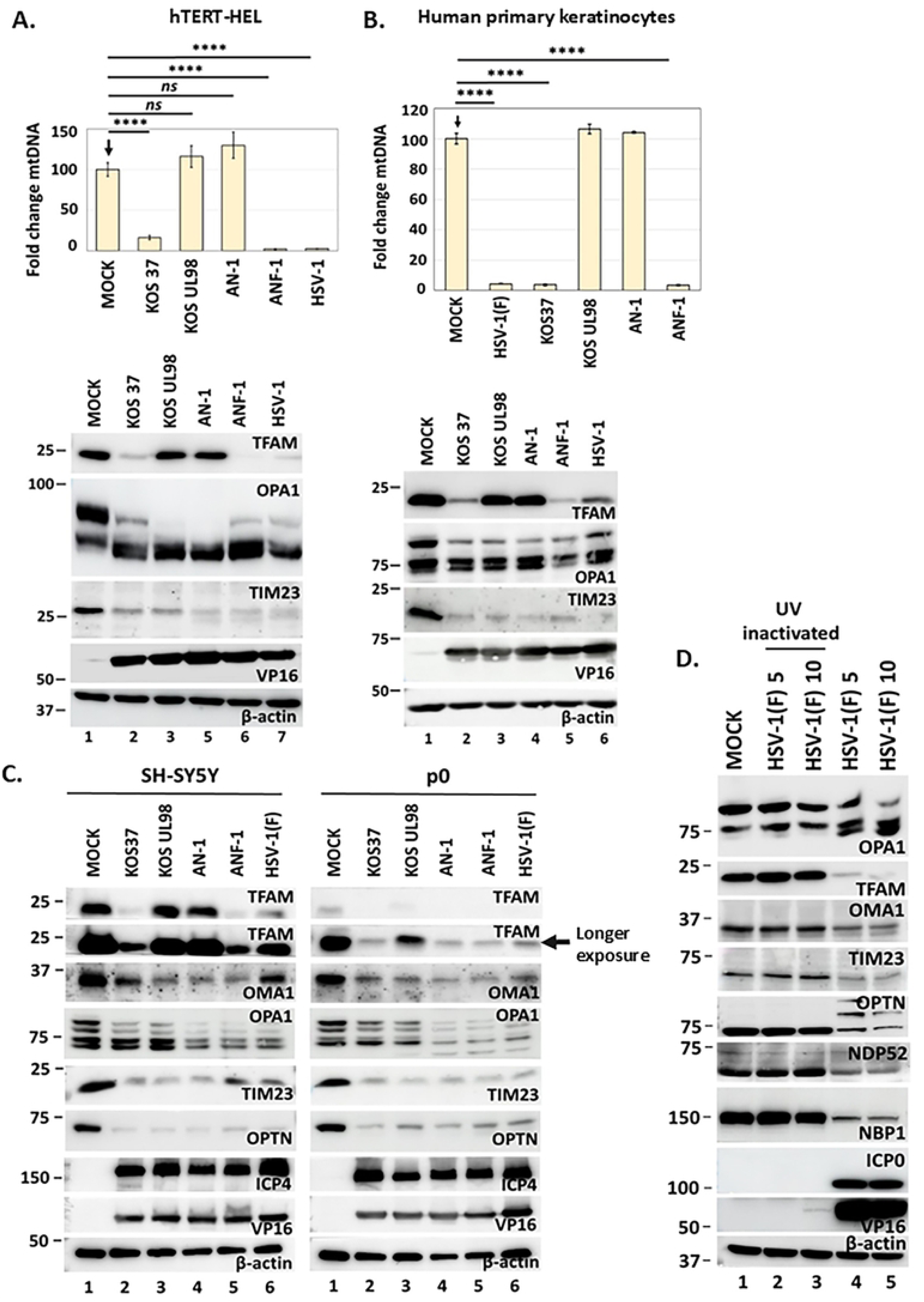
UL12.5 causes mtDNA/TFAM degradation in HSV-1–infected cells. **A.** hTERT-HEL were infected with KOS37, KOS UL98, AN-1, ANF-1, and HSV-1(F) (5 PFU/cell) or left uninfected. The cells were harvested at 14 h post-infection and equal amounts of total DNA were analyzed for mtDNA by qPCR. Nuclear DNA was used for normalization. Cells were also harvested at 14 hpi and equal amounts of proteins were analyzed for TFAM, OPA1, TIM23, VP16, and β-actin. **B.** Human primary epidermal keratinocytes were infected as in panels A. Analysis of protein and mtDNA was performed at 14 h post-infection as above. **C.** SH-SY5Y and ρ0 derivatives were infected with HSV-1(F) (5.0 PFU/cell) or left uninfected. The cells were harvested at 18 h post-infection and equal amounts of proteins from total cell lysates were analyzed for TFAM (a shorter and longer exposure are depicted), OMA1, and TIM23. ICP4 and VP16 served as controls for the infection, and β-actin as a loading control. **D.** hTERT-HEL cells were infected with two doses of HSV-1(F) or UV-inactivated virus (5 and 10 PFU/cell) or left untreated. The cells were harvested at 14 h post-infection and equal amounts of proteins from total cell lysates were analyzed for OPA1, TFAM, OMA1, TIM23, OPTN, NDP52, and NBR1. ICP0 and VP16 served as controls for the infection as well as to determine the efficiency of UV inactivation, and β-actin as a loading control. For all the experiments statistical significance was calculated from three biological replicates with two-tailed Student’s *t*-test. P<0.0001****, ns=not significant.

Next, we assessed whether loss of TFAM in HSV-1-infected cells depends on mtDNA. For this we utilized a neuroblastoma cell line (SH-SY5Y) that was depleted of mtDNA by ethidium bromide treatment (ρ0) (68–70). The ρ0 cell line expresses TFAM, but the protein does not accumulate to high levels, due to the absence of mtDNA that is required for TFAM protein stabilization. SH-SY5Y cells with (p+) and without (ρ0) mtDNA were infected with HSV-1(F), KOS37, AN-1, ANF-1, and KOS UL98 (5 PFU/cell), and we assessed TFAM protein levels. We discovered that TFAM was depleted in both cell lines after infection with the UL12.5-expressing viruses (KOS37, ANF-1, and HSV-1(F)) (Figure 5C). However, in the p0 cell line, TFAM was also depleted in the absence of UL12.5 or when HCMV UL98 was expressed. These data indicate that in the absence of mtDNA, a UL12.5–independent mechanism degrades TFAM during HSV-1 infection. These findings also show that HSV-1 can initiate both UL12.5–dependent and UL12.5–independent signals for TFAM depletion, depending on the environment of infection. We also observed that OMA1 activation and degradation of TIM23 occurred in a UL12.5–independent manner in both cell lines. Viral gene expression was comparable between the two cell lines, indicating that mtDNA is not required for optimal viral gene expression (Figure S3C). We also asked whether virus entry to cells is sufficient to cause some of the mitochondrial changes described earlier. hTERT-HEL cells were infected with two doses of HSV-1(F) (5 and 10 PFU/cell), and equal dose of UV-inactivated virus. The cells were harvested at 14 h post-infection and analyzed for OPA1 conversion, OMA1 activation, TFAM and TIM23 depletion, as well as any effects on the mitophagy adaptor proteins OPTN (optineurin), NDP52 (nuclear dot protein 52 kDa), and NBR1 (Neighbor of BRCA gene 1) (Figure 5D). We found that virus entry to the cells was not sufficient to cause the mitochondrial changes described before. Altogether, these data indicate that expression of UL12.5 is required for mtDNA/TFAM loss during HSV-1 infection. UL12.5–independent signals inhibit mitochondrial fusion and cause TIM23 down-modulation, but these signals are not sufficient to cause mtDNA/TFAM loss.

Additionally, we asked whether UL12.5 alone is sufficient for the mitochondrial changes discussed earlier. UL12.5-EGFP was abundantly present in mitochondria both in infected and uninfected cells (Figure 6A). We developed a doxycycline (Dox)-inducible UL12.5 hTERT-HEL cell line and confirmed that UL12.5 localizes to mitochondria both when expressed alone and in the context of infection (data not shown). We then found that UL12.5 alone was sufficient to cause TFAM and mtDNA loss, but did not cause OMA1 activation, OPA1 conversion, or TIM23 loss (Figure 6B). Further, we found that inhibiting lysosomal activity with bafilomycin A1 (BafA1) early upon UL12.5 expression, resulted in an increased accumulation of a shorter UL12.5 protein (likely after MLS cleavage or a UL12-M185 form, previously reported (24)) as well as a partial rescue of TFAM and mtDNA (Figure 6C, and data not shown). This is consistent with previous studies proposing the disposal of enlarged mt-nucleoids into lysosomes (27). The LonP1 inhibitor did not restore TFAM and mtDNA levels in the UL12.5-expressing cell line (data not shown). Overall, UL12.5 alone is sufficient to cause TFAM and mtDNA loss.

**Figure 6:**
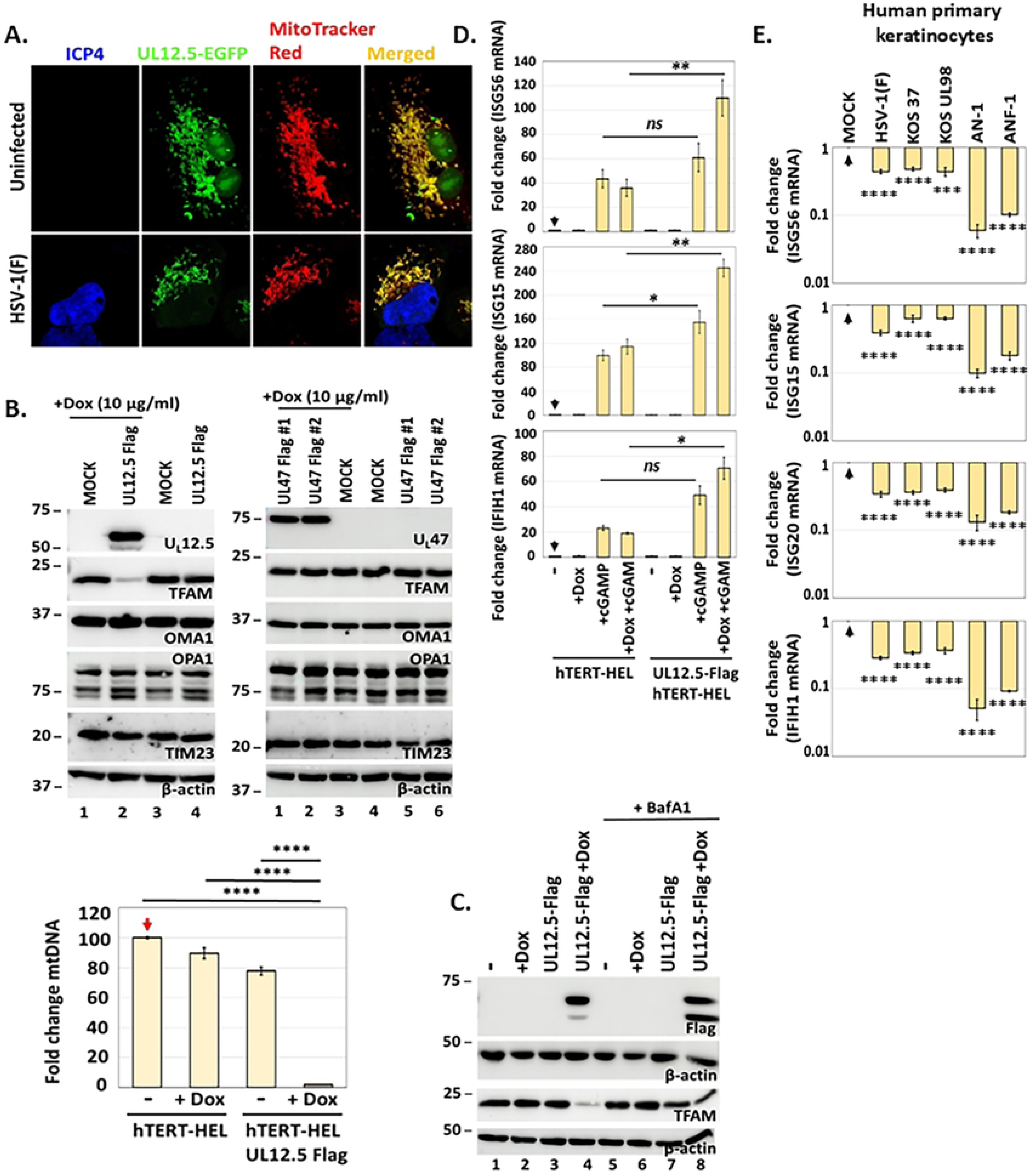
UL12.5 alone is sufficient to cause TFAM and mtDNA degradation. **A.** ARPE-19 cells were transfected with a UL12.5-EGFP-expressing plasmid and infected with HSV-1(F) (10 PFU/cell) or left uninfected. Live cells were stained with MitoTracker Red, fixed at 14 h post-infection, and images were acquired using a Leica TCS SP8 confocal microscope. **B.** An inducible UL12.5-Flag hTERT-HEL cell line and two UL47-Flag cell lines were treated with Dox (10 μg/ml) for 24 h or left untreated. Dox was removed to avoid potential stress on mitochondria, samples were harvested at 72 h post-treatment, and equal amounts of proteins from total cell lysates were analyzed for UL12.5, UL47, TFAM, OMA1, TIM23, and β-actin. The levels of mtDNA were quantified from the UL12.5-expressing samples, as described earlier. **C.** The UL12.5-Flag hTERT-HEL cell line was treated with Dox (10 μg/ml) and BafA1 (15 μM) simultaneously or left untreated. Samples were harvested at 12 h post-treatment and equal amounts of proteins from total cell lysates were analyzed for UL12.5 and TFAM. As a loading control we used β-actin. **D.** The UL12.5-Flag hTERT-HEL cell line was treated with Dox (10 μg/ml) for 4 h or left untreated. Then, 2’3’-cGAMP (12 μM) was added to the cultures for 14 h. Total RNA was extracted from three biological replicates and analyzed for ISGs expression by RT-qPCR. **E.** Human primary epidermal keratinocytes were infected with HSV-1(F), KOS37, KOS UL98, AN-1, and ANF-1 virus (0.1 PFU/cell). The cells were harvested at 14 h post-infection, total RNA was extracted, and quantification of *ISG56*, *ISG15*, *ISG20*, and *IFIH1* transcripts was done by RT-qPCR. Normalization was done against 18S rRNA. For all the experiments statistical significance was calculated from three biological replicates with two-tailed Student’s *t*-test. P<0.05*, P<0.01**, P<0.001***, P<0.0001****, ns= not significant.

Finally, previous studies indicated that disruption of mtDNA by UL12.5 can activate type I IFN signaling (52). Therefore, we tested whether UL12.5 expression or infection with the UL12/UL12.5 mutant viruses cause type I IFN gene expression. We found that UL12.5 expression alone did not stimulate expression of the interferon stimulated genes (ISGs) -*ISG56*, *ISG15*, and *IFIH1-* in hTERT-HEL cells (Figure 6D). However, treatment of the UL12.5-expressing cells with the non-canonical cyclin dinucleotide, 2’3’-cGAMP, stimulated higher expression of type I IFN genes compared to cells that did not express UL12.5. These findings indicate that UL12.5 plays a role in priming the cells for activation of cell defense signaling. None of the UL12.5 mutant viruses tested activated type I IFN responses in human primary epidermal keratinocytes, hTERT-HEL, or HaCaT cells where the virus productively replicates (Figures 6E, S4A, and data not shown). This is likely because all of the viral genes required for immune evasion are expressed by these mutant viruses (71;72). However, we did notice a marginal increase of LC3 lipidation after infection with the AN-1 virus (UL12/UL12.5 null mutant) (Figure S4B), perhaps because of an accumulation of defective particles within AN-1-infected cells (72). Taken together, our findings reveal both UL12.5-dependent and UL12.5-independent mechanisms that contribute to the disruption of mitochondrial homeostasis in HSV-1-infected cells.

### Mitophagy is inhibited in HSV-1 infected cells to preserve the mitochondrial network

Due to extensive mitochondrial changes in HSV-1–infected cells, we investigated whether mitophagy could clear those mitochondria. We previously reported that two major autophagy/mitophagy adaptors, p62/SQSTM1 (sequestosome1) and OPTN, are down-modulated in HSV-1 infected cells via an ICP0-dependent mechanism (39). Here, we investigated whether additional autophagy/mitophagy adaptors are affected in HSV-1–infected cells. We performed kinetic studies in hTERT-HEL cells infected with HSV-1(F) and determined that NBR1 and NDP52 are also down-modulated, like p62/SQSTM1 and OPTN (Figures 7A, 7C). Similar results were obtained in human primary epidermal keratinocytes, ARPE-19, and HaCaT (data not shown). A mitophagy pathway that is frequently activated upon stress involves PINK1, which accumulates to the OMM and recruits the E3 ubiquitin ligase, Parkin. Subsequently, Parkin ubiquitinates mitochondrial proteins marking them for clearance through mitophagy (15). PINK1/Parkin are upregulated during the early stages of mitophagy and depleted through the process (15). We monitored levels of PINK1 and Parkin during infection with the WT and ΔUL47 virus (2.5 PFU/cell) and observed only a marginal decrease of the mature and dimeric PINK1 at 24 h post-infection, with no effect on Parkin (Figure 7B). These data indicate that HSV-1 infection does not activate mitophagy. We built upon these findings and tested whether the expression of the autophagy/mitophagy adaptors could be rescued in cells depleted of ATG5 (autophagy-related protein 5), where autophagosome formation is blocked. We found that ATG5 depletion could not rescue these adaptor proteins (Figure 7C). As a positive control, we monitored LC3 (microtubule-associated protein 1 light chain 3). LC3 is a key structural component of autophagosomes. Upon activation of autophagy, LC3 becomes lipidated and, as a result, migrates faster on a denaturing gel. We found that ΔUL47 virus could not efficiently evade autophagy in hTERT-HEL cells, which caused accumulation of lipidated LC3 (Figure 7C). LC3 lipidation was not observed in ΔUL47 virus –infected ATG5 KD cells, indicating that autophagy was efficiently blocked in this cell line (Figure 7C). We also tested if mtDNA could be rescued in HSV-1–infected ATG5 KD cells but found that this was not the case either (Figure 7D). These data support the conclusion that autophagy/mitophagy is not responsible for mtDNA down-modulation.

**Figure 7:**
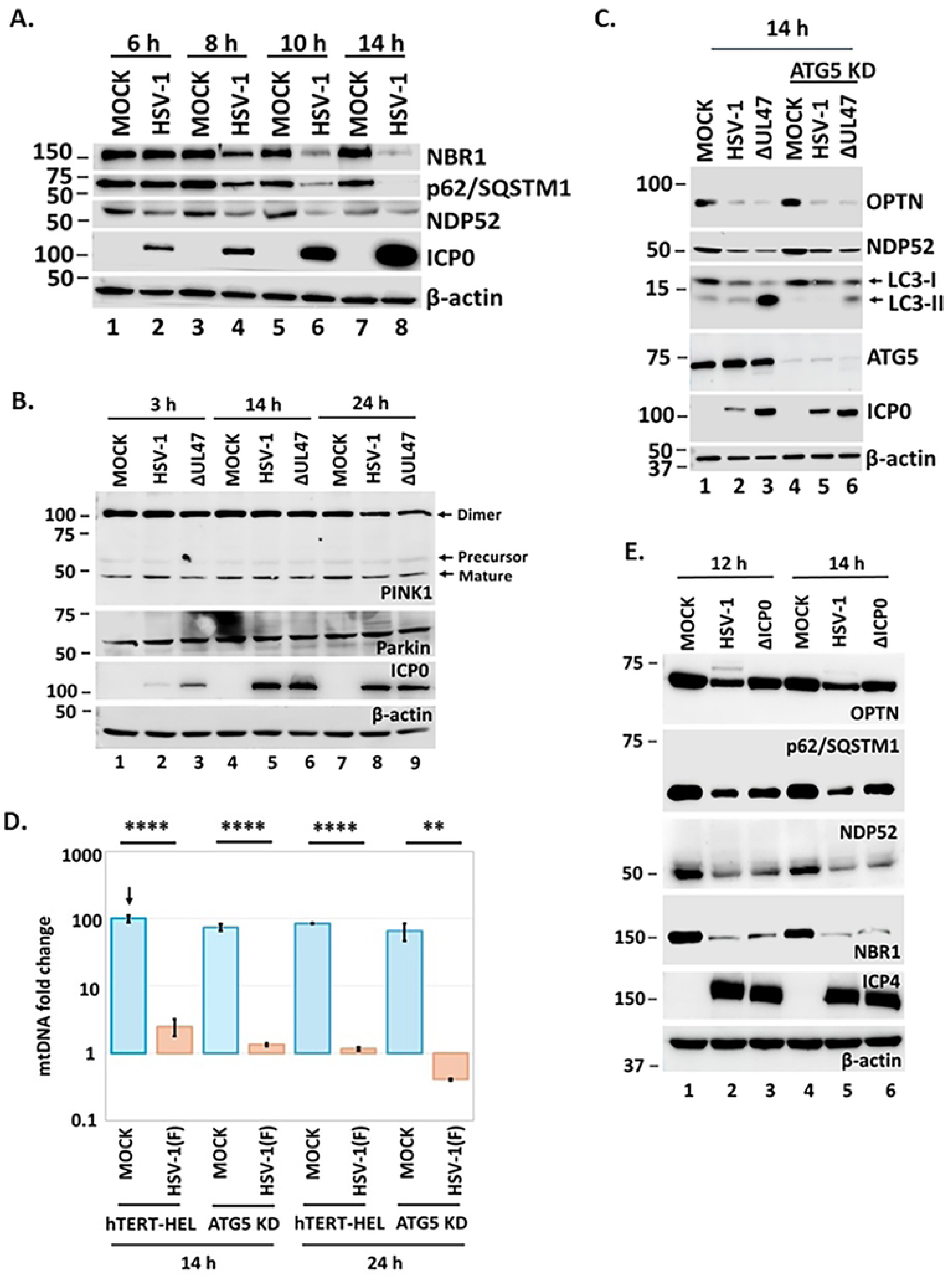
HSV-1 infection down-modulates several mitophagy adaptor proteins. **A.** hTERT-HEL cells were infected with HSV-1(F) (5 PFU/cell). The cells were harvested at 6, 8, 10, and 14 h post-infection and equal amounts of proteins from total cell lysates were analyzed for NBR1, p62/SQSTM1, and NDP52. ICP0 served as a control for infection and β-actin as a loading control. **B.** hTERT-HEL cells were infected with HSV-1(F) and ΔUL47 virus (2.5 PFU/cell). The cells were harvested at 3, 14, and 24 h post-infection and equal amounts of proteins from total cell lysates were analyzed for PINK1 and Parkin. ICP0 served as a control for infection and β-actin as a loading control. **C.** hTERT-HEL cells and ATG5 KD derivatives were infected with HSV-1(F) and ΔUL47 virus (2.5 PFU/cell). The cells were harvested at 14 h post-infection and equal amounts of proteins from total cell lysates were analyzed for OPTN, NDP52, ATG5, and LC3. ICP0 served as a control for infection and β-actin as a loading control. The faster migrating form of LC3 represents a lipidated form. **D.** hTERT-HEL cells and ATG5 KD derivatives were infected with HSV-1(F) (2.5 PFU/cell). The cells were harvested at 14 and 24 h post-infection, total DNA was extracted and mtDNA was quantified by qPCR. Results from three independent experiments are depicted. Statistical significance was calculated with two-tailed Student’s *t*-test. P<0.01**, P<0.0001****. **E.** hTERT-HEL cells were infected with HSV-1(F) and ΔICP0 virus (5 PFU/cell). The cells were harvested at 12 and 14 h post-infection and equal amounts of proteins from total cell lysates were analyzed for OPTN, p62/SQSTM1, NDP52, and NBR1. ICP4 served as a control for infection and β-actin as a loading control.

Previously, we described that p62/SQSTM1 and OPTN are down-modulated in an ICP0-dependent manner. However, this was not the case for NDP52 and NBR1 (Figure 7E). For this reason, we investigated whether the lysosomal inhibitor BafA1, or the proteasome inhibitor MG132, could rescue the proteins that we previously found to be eliminated. Both drugs were added at 4 and at 6 h post-infection to avoid interference with viral gene expression, and proteins were analyzed at 14 h post-infection (Figure S4C). We determined that BafA1 could partially rescue TFAM, but neither BafA1 nor MG132 could rescue TIM23, NBR1, MFN2, or OMA1 (Figure S4C). These data indicate that lysosomal activity has only a minor role in eliminating mitochondrial-associated factors during HSV-1 infection.

Overall, these data indicate that mitophagy is inhibited in HSV-1-infected cells and that mtDNA degradation occurs in a mitophagy-independent manner. Lysosomes likely have a contributing role in removing some mitochondrial content.

### HSV-1 infection causes exocytosis of mtDNA and selected mitochondrial factors

Recent studies have indicated that enlarged mt-nucleoids can enter endosomal pathways and traffic to lysosomes for degradation (27). Thus, we investigated a potential interplay between endosomes and mitochondria in HSV-1–infected cells. To visualize this, we co-expressed the DNA helicase Twinkle, which binds to mtDNA, with early (Rab5) and late (Rab7, LAMP1) endosomal markers in ARPE-19 cells (human retinal epithelial). The transfected cells were infected with HSV-1(F) (10 PFU/cell) or left uninfected, the samples were fixed at 16 h post-infection, and images were acquired with the Leica TCS SP8 confocal microscope. We discovered that both early and late endosomal markers were proximal to Twinkle (diffraction limit of confocal microscopy is > 200 nm). In infected cells these proximity events frequently resulted in partial overlap of Twinkle with Rab5, Rab7, and LAMP1 (Figure S5A-B). However, in uninfected cells these overlapping events were less frequent. We obtained similar results in cells co-transfected with TOM20 (an OMM protein) and the same endosomal markers (Figure S5C and data not shown). These results indicate that endosomes and mitochondria are likely exchanging content with higher frequency during HSV-1 infection.

Considering the proximity of endosomes to mitochondria during HSV-1 infection, the failure of BafA1 to restore mitochondrial protein levels, and the inhibition of mitophagy altogether indicated that mitochondrial factors, which enter an endosomal pathway, may be targeted for exocytosis instead of autophagolysosomal degradation. To test this, hTERT-HEL cells were infected with HSV-1(F) (0.5 PFU/cell) or left uninfected. Culture supernatants were collected at 24 h post-infection and EVs were isolated and analyzed for selected mitochondrial factors (Figure 8A). TFAM, whose levels decreased intracellularly in HSV-1(F) infected cells, was detected in EVs (Figure 8A), but this was not the case for TIM23. The OMM proteins TOM20, TOM40, and VDAC1 were also present in EVs from infected cells, but not TOM70 or MFN2 (Figure 8A). None of the IMM proteins (TIM23, TIM44, or OPA1) were present in EVs. We also looked at nuclear-encoded ETC components and found that UQCRC1, but not NDUFA9 was present in EVs (Figure 8A). C1QBP, a multifunctional protein with roles in mitochondrial regulation and functions, was also found in EVs. Finally, early (Rab5) and late endosomal markers (Rab7 and LAMP1) that intercalate with mitochondria, the SNARE protein Syntaxin 17 (facilitates fusion of autophagosomes with lysosomes), and mitophagy adaptor proteins, were also present in EVs from infected cells (Figure 8A). Since Alix, an accessory protein of the ESCRT (endosomal sorting complexes required for transport) machinery, does not change significantly in lysates and EVs between infected and uninfected cells, it was used as a loading control (73;74). Overall, we observed that selected OMM proteins, along with nuclear-encoded ETC factors and mitophagy adaptors, are exocytosed during HSV-1 infection.

**Figure 8:**
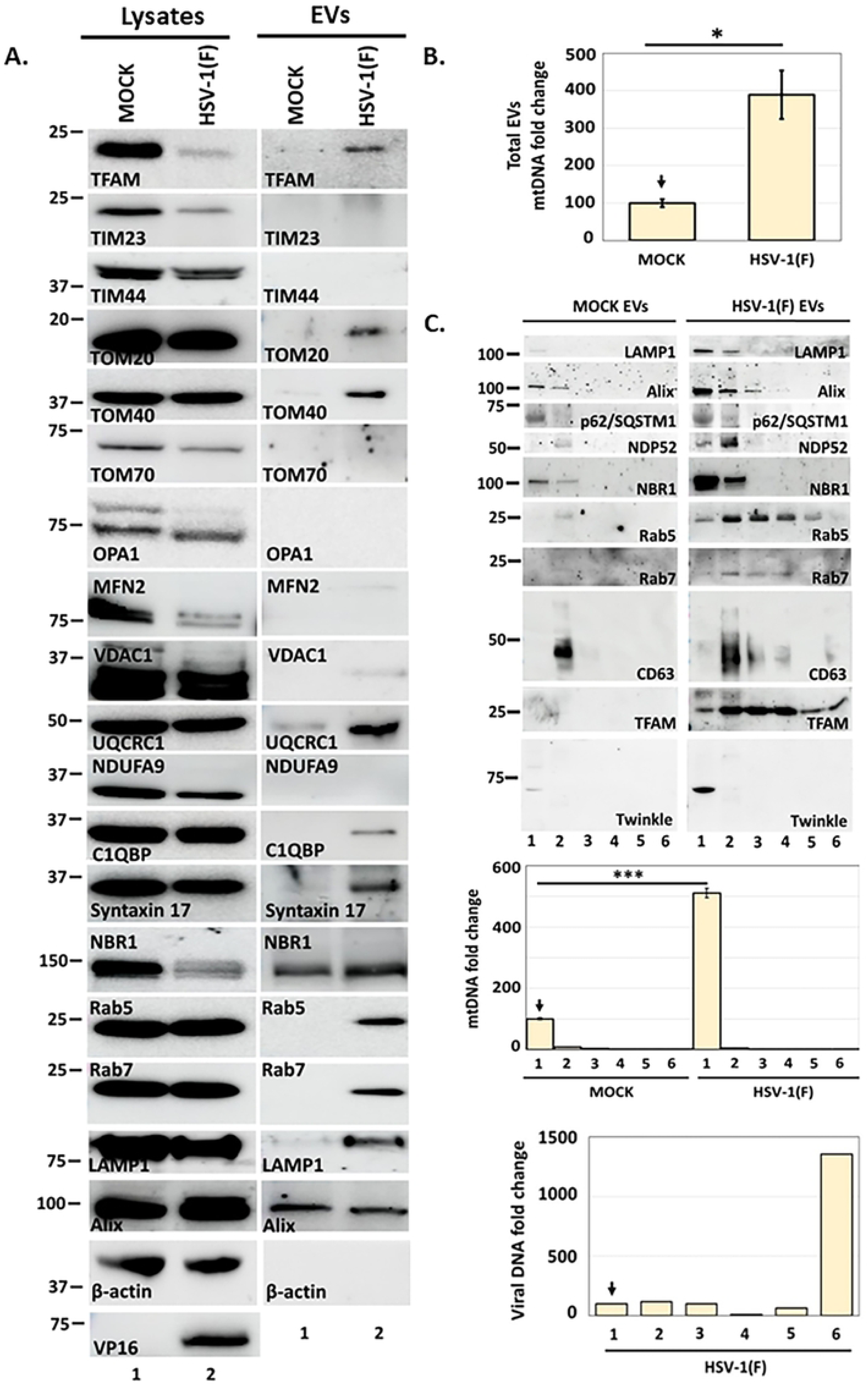
mtDNA and other mitochondrial factors are present in EVs released by HSV-1(F) infected cells. **A.** hTERT-HEL cells were infected with HSV-1(F) (0.5 PFU/cell) or left uninfected. Culture supernatants were harvested at 24 h post-infection, clarified in low-speed centrifugation, concentrated using 100 kDa cutoff filters, and pelleted at 100,000x *g*. Equal amounts of proteins from pelleted EVs and cell lysates were analyzed for TFAM, TIM23, TIM44, TOM20, TOM40, TOM70, OPA1, MFN2, VDAC1, UQCRC1, NDUFA9, C1QBP, Syntaxin 17, NBR1, Rab5, Rab7, and LAMP1. VP16 served as a control for the infection, β-actin as a loading control of cell lysates, and Alix as a loading control for EVs. **B.** hTERT-HEL cells were infected with HSV-1(F) (0.5 PFU/cell) or left uninfected. Culture supernatants were harvested at 24 h post-infection, clarified at low-speed centrifugation, concentrated using 100 kDa cutoff filters, spun at 15,000x *g* to remove virions and larger vesicles, the supernatant containing EVs was further concentrated through 100 kDa cutoff filters, total DNA was extracted, and mtDNA was quantified by qPCR using equal volume of total DNA. Statistical significance was evaluated using three replicates with two-tailed Student’s *t*-test. P<0.05*. **C.** hTERT-HEL cells were infected with HSV-1(F) (0.5 PFU/cell) or left uninfected. Culture supernatants were harvested at 24 h post-infection, subjected to low-speed centrifugation and concentrated, as above. Concentrated culture supernatant was subjected to a centrifugation at 15,000x *g* to remove virions and larger vesicles, the supernatant was further concentrated through 100 kDa cutoff filters, loaded on a 6-18% iodoxanol/sucrose gradient, and subjected to 250,000x *g* centrifugation. Two milliliter fractions were collected from the top to the bottom of the gradient and analyzed for different proteins. Also, mtDNA was isolated and quantified by qPCR in each fraction. Viral DNA was also quantified in the same fractions by qPCR using primers targeting the gI locus.

Next, we asked if mtDNA is present in EVs released by HSV-1-infected cells. To test this, we first analyzed the total EV content released by HSV-1 infected cells for the presence of mtDNA. hTERT-HEL cells were infected with HSV-1(F) (0.5 PFU/cell) or left uninfected. Culture supernatants were harvested at 24 hpi, clarified at low-speed centrifugation, filtered/concentrated, and subjected to a 15,000x *g* centrifugation to remove larger vesicles and virus. The EVs in the supernatant were then concentrated through 100 kDa cutoff filters and mtDNA was quantified by qPCR. We discovered that infected cells release approximately 4-fold more mtDNA compared to uninfected cells (Figure 8B). However, this is a crude process that cannot separate EVs from virions. Therefore, we implemented a gradient approach that we previously developed to separate EVs from virions and identified the fractions of the gradient containing mitochondrial proteins and mtDNA(73;75-77). Culture supernatants were harvested at 24 hpi, clarified at low-speed centrifugation, filtered/concentrated through 100 kDa cutoff filters, and subjected to a 15,000x *g* centrifugation to remove larger vesicles and virions. These populations were then further concentrated through 100 kDa cutoff filters or pelleted at 100,000x *g*, loaded on a 6-18 % iodixanol/sucrose gradient, and subjected to a 250,000**x** *g* ultracentrifugation. Two milliliter fractions were then collected from the top to the bottom of the gradient for protein and mtDNA analysis. As shown in Figure 8C, approximately 5-fold more mtDNA was detected in the first fraction of the gradient from infected compared to uninfected cells. In the same fraction, we detected TFAM and Twinkle (the mtDNA helicase), indicating that mtDNA and its bound proteins are both exocytosed, and that they co-fractionate. We also detected selected endosomal markers (LAMP1, Alix, Rab5 and Rab7), and the mitophagy adaptors (NBR1 and NDP52) that are potentially involved in mtDNA/TFAM/Twinkle exocytosis (Figure 8C). Notably, the tetraspanin CD63 whose exocytosis is induced in HSV-1(F) infected cells did not co-fractionate with mtDNA, indicating that mtDNA is not exocytosed in CD63+ EVs (73;76;77). To verify the purity of EVs, we also probed for viral DNA and found that it accumulates in the bottom fraction of the gradient (fraction 6) (Figure 8C).

Taken together, in HSV-1–infected cells, we show that mtDNA, along with mtDNA–binding proteins, and several OMM proteins, enter an endosomal pathway and are sorted for exocytosis. We also observe that a small fraction of this cargo may be targeted to lysosomes for degradation.

### HSV-1 infection increases spare respiratory capacity and the extracellular acidification rate, and does not disrupt mitochondrial membrane potential

Here, we analyzed functional aspects of mitochondria in HSV-1 infected cells. It has been reported that mitochondria in HSV-1-infected cells accumulate to perinuclear sites, likely to supply energy and metabolites for virus replication and virion assembly. We verified these findings in infected hTERT-HEL cells constitutively expressing mitochondrial targeted EGFP (OMP25-EGFP), and in infected hTERT-HEL stained with MitoTracker Red, or an OPA1 antibody (Figure 9A). In all cases, we noticed a perinuclear accumulation of mitochondria. These mitochondria co-clustered with virion assembly sites that were identified by staining with an anti-HSV-1 antibody raised against virion components (Figure 9A). Despite these changes, mitochondria from HSV-1–infected cells are likely functional, and this is probably the reason why they cluster at the virus assembly sites. Similar perinuclear clustering of mitochondria has been observed when there is an increased energy demand in the nucleus (78).

**Figure 9:**
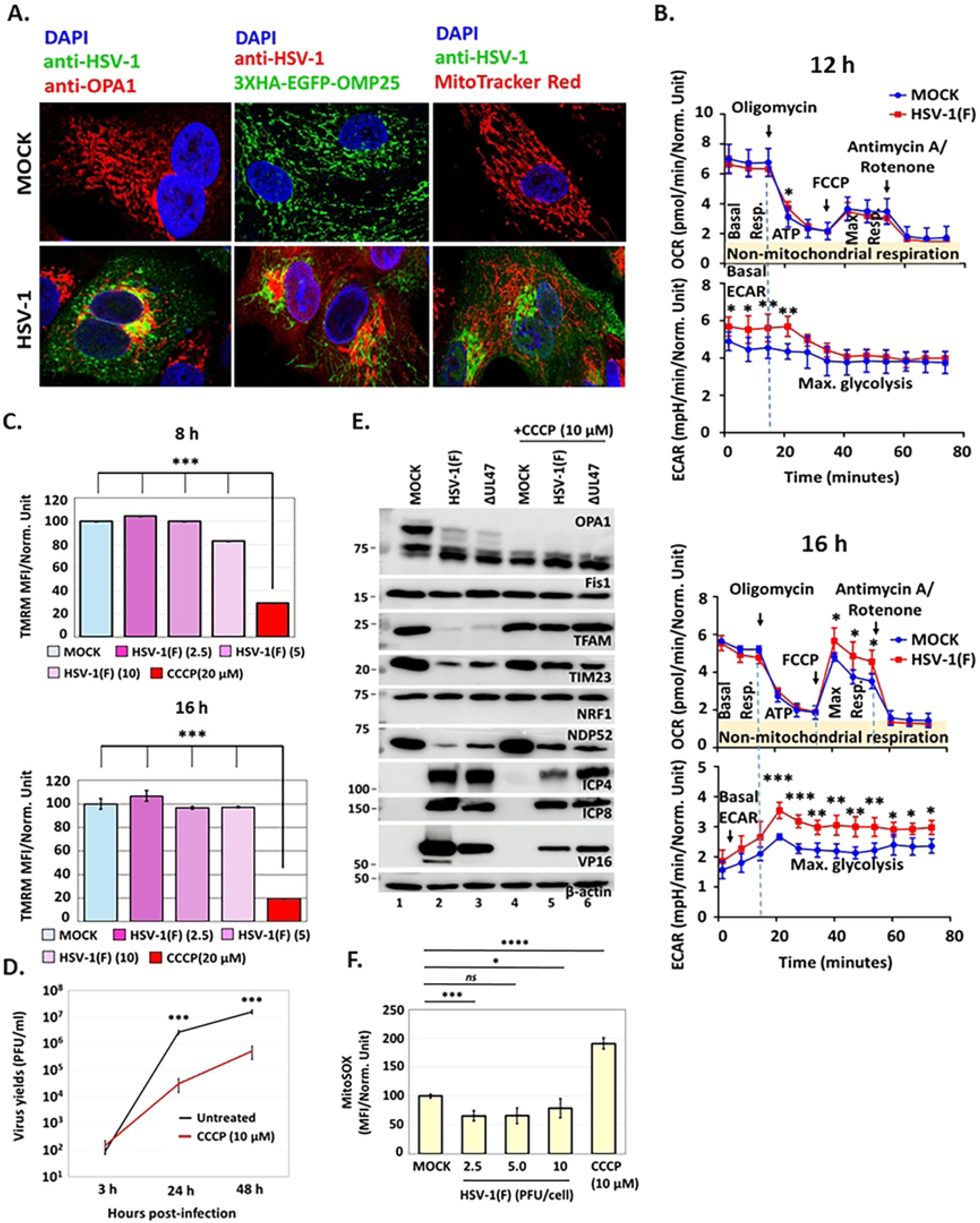
Mitochondria maintain certain metabolic activity in HSV-1 (F) infected cells. **A.** Replicate cultures of hTERT-HEL cells were infected with HSV-1(F) (1 PFU/cell) or left uninfected. The samples were fixed at 16 h post-infection and doubly stained with an anti-OPA1 and anti-HSV-1/2 antibody. In addition, an hTERT-HEL cell line expressing mitochondrial targeted EGFP (EGFP-OMP25) was infected with HSV-1(F) (1 PFU/cell), fixed at 16 hpi, and stained with an anti-HSV-1/2 antibody. Further, hTERT-HEL cells were infected with HSV-1(F) (1 PFU/cell) and incubated with MitoTracker Red. The samples were fixed at 16 hpi and probed with an anti-HSV-1/2 antibody. Images were acquired with a Leica TCS SP8 confocal microscope. **B.** hTERT**-**HEL cells were infected with HSV-1(F) (2 PFU/cell) or remained uninfected. At 12 and 16 hpi a Seahorse analysis was performed to monitor OCR and ECAR. Oligomycin (2 μM), FCCP (1 μM), and antimycin A (1 μM)/rotenone (50 nM) were used at the indicated time and concentrations. Results were normalized to the number of cells (1,000 cells). Statistical significance was calculated from eight biological replicates with two-tailed Student’s *t*-test. P<0.05*, P<0.01**, P<0.001***. **C.** hTERT-HEL cells were infected with HSV-1(F) (2.5, 5, 10 PFU/cell). CCCP (20 μM) was added to the cultures at 3 hpi, cells were stained with TMRM, and live cell imaging was performed at 8 and 16 hpi in Cytation 7. Mean fluorescence intensity (MFI) of the cells was calculated using the Gen5 Image+ 3.13 software. Statistical significance was calculated using three biological replicates and two images per replicate, with two-tailed Student’s *t*-test. P<0.05*, P<0.01**, P<0.001***. **D.** hTERT-HEL cells were infected with HSV-1(F) (0.01 PFU/cell). CCCP (10 μΜ) was added to the infected cultures at 3 hpi. The cells were harvested at 3, 24, and 48 hpi and progeny virus was quantified by plaque assays in Vero cells. **E.** hTERT-HEL cells were infected with HSV-1(F) and ΔUL47 virus (2.5 PFU/cell). CCCP (10 μM) was added to the infected cultures at 3 hpi. The cells were harvested at 24 hpi and equal amounts of proteins from total cell lysates were analyzed for OPA1, Fis1, TFAM, TIM23, NRF1, and NDP52. ICP4, ICP8, and VP16 were used to monitor the infection, and β-actin as a loading control. **F.** hTERT-HEL cells were infected with HSV-1(F) (2.5, 5, 10 PFU/cell) or left uninfected. MitoSOX (50 μM) was added to the live cultures for 45 min. Images were acquired at 16 hpi using the Gen5 Image+ 3.13 software of Cytation 7. Replicate cultures treated with CCCP (10 μM) served as a positive control. Statistical significance for all experiments was calculated from at least three biological replicates with two-tailed Student’s *t*-test. P<0.05*, P<0.01**, P<0.001***.

ATP in mammalian cells is generated by mitochondrial (oxidative phosphorylation) and non-mitochondrial (glycolysis) metabolism. To monitor mitochondrial metabolic activity in HSV-1(F)-infected cells we measured the oxygen consumption rate (OCR) and the extracellular acidification rate (ECAR). OCR measures respiration and spare respiratory capacity, whereas ECAR measures glycolysis and proton flux (acidification). Measurement of OCR and ECAR was done simultaneously on live cultures using a Seahorse XF Pro (Figure 9B). Key parameters of respiration (basal respiration, ATP-linked respiration, maximal respiratory capacity, reserved respiratory capacity, non-mitochondrial respiration) and glycolysis (basal ECAR, maximal glycolysis) were assessed in hTERT-HEL infected with HSV-1(F) (2 PFU/cell) versus uninfected cells, at 12 and 16 hpi, by sequentially exposing cells to mitochondrial perturbing reagents. After establishing a baseline for basal OCR and ECAR, oligomycin (20 μM) was added to the cultures to inhibit ATP synthase. This typically causes a decrease in OCR and an increase in ECAR because the ETC is blocked from producing ATP, which forces the cells to rely on glycolysis, leading to extracellular acidification. Following oligomycin treatment we did not notice any difference in OCR between infected and uninfected cells, but we did notice a statistically significant increase in ECAR during infection at both time points. FCCP (carbonyl cyanide 4-(trifluoromethoxy) phenylhydrazone) (1 μM) followed oligomycin treatment. FCCP uncouples the ETC from ATP production, causing a rapid increase in OCR and a subsequent increase in glycolysis (and ECAR), due to reduced ATP generation within the mitochondria. We noticed a statistically significant increase in both OCR and ECAR in infected cells at 16 hpi (Figure 9B). Finally, the cultures were injected with antimycin A (20 nM) and rotenone (8 nM), two ETC inhibitors, which decreased the OCR, shutting down respiration. OCR was comparable between infected and uninfected cells, whereas ECAR remained at higher levels in infected compared to uninfected cells at 16 hpi (Figure 9B). Altogether, these data indicate that HSV-1 infection increases mitochondrial spare respiration and glycolytic activity over time, in order to support the infection.

We then assessed whether HSV-1(F) disrupts the mitochondrial membrane potential (Δψm), which is crucial for maintenance of mitochondrial integrity and function. hTERT-HEL cells were infected with HSV-1(F) at different doses (2.5, 5.0, and 10 PFU/cell), and stained at 8 and 16 hpi with TMRM (tetramethylrhodamine, methyl ester), a cell-permeant dye that accumulates in mitochondria with intact membrane potential. The fluorescent signal was quantified with the Gen5 software of Cytation 7. We discovered that HSV-1(F) infection does not disrupt Δψm (Figure 9C). As a positive control we treated the cells with CCCP (20 μM), which caused Δψm loss and a decreased TMRM signal.

To determine how Δψm loss could impact the infection we infected hTERT-HEL cells with HSV-1(F) (0.01 PFU/cell), treated the cells with CCCP (10 μM) at 3 hpi, and analyzed progeny virus production at 24 and 48 hpi. We found that CCCP treatment decreased progeny virus production by at least 2-log (Figure 9D). CCCP treatment delayed all classes of viral genes, and this, in turn, decreased the ability of the virus to eliminate TFAM and TIM23 (Figure 9E). However, the virus could still down-modulate the autophagy/mitophagy adaptors (such as NDP52), as this is an earlier event. CCCP treatment promoted mitochondrial fission and the extensive conversion of L-OPA1 to S-OPA1 (Figure 9E). Notably, HSV-1 infection causes L-OPA1 to S-OPA1 conversion to a similar extent as CCCP treatment, which further supports that HSV-1 infection blocks fusion. Finally, we monitored oxidation of the superoxide indicator MitoSOX Red. This is a fluorogenic dye that specifically targets mitochondria in live cells and upon oxidation produces red fluorescence. We found that MitoSOX Red signal was comparable between infected and uninfected cells (Figure 9F). These data indicate that HSV-1 infection does not induce mitochondrial superoxide production. As a positive control, CCCP was used at a high concentration (10 μM), since it is known to increase reactive oxygen species (ROS) production (Figure 9F).

Overall, despite all the mitochondrial membrane changes mitochondria retain certain metabolic activity during HSV-1 infection.

## Discussion

The role of UL12.5 in mtDNA degradation during HSV-1 infection has been puzzling. UL12.5 protein is an N-terminal truncated form of the UL12 alkaline deoxyribonuclease. UL12 functions in the nucleus during HSV-1 replication to resolve branched concatemeric DNA structures to linear unit-length genomes that can be packaged into the capsids (79). To perform this function, UL12 has both endo- and exo-nuclease activities and functions as a two-component recombinase with the single-stranded viral DNA-binding protein ICP8 (79). UL12 also interacts with DNA repair proteins for efficient virus replication (80). UL12.5 retains these functions but localizes to mitochondria (24;26;63-65). The weak nuclease activity of UL12.5 is dispensable for mtDNA/TFAM degradation, indicating that another UL12.5 function is responsible (26). Instead, it has been proposed that mitochondrial endonucleases may play redundant roles in UL12.5-mediated mtDNA depletion (26). Mitochondria contain only a small number of endonucleases, most of which act on specialized DNA substrates (81). An exception is the endonuclease G (ENDOG), which shares similar biochemical properties with UL12.5, such as the requirement for magnesium, alkaline pH, and low ionic strength, and has been implicated in mtDNA degradation during HSV-1 infection (81). Similarly, exonuclease G (EXOG) has been implicated in mtDNA degradation during HSV-1 infection (26). EXOG has both endonuclease and weak 5΄→3΄ exonuclease activity with preference for single-stranded DNA (81). Despite these findings, depletion of both endonucleases did not fully rescue mtDNA levels (26). Also, direct interaction of UL12.5 with these nucleases has not been demonstrated. These data indicate that another mechanism mediates mtDNA degradation through UL12.5. A recent study proposed that ectopic expression of UL12.5 results in enlarged mt-nucleoids (27). Enlarged mt-nucleoids are an indication of mitochondrial dysfunction, a replication and/or fission defect, or the presence of a stressor such as a virus. However, these studies did not propose any specific function for UL12.5.

Our studies provide mechanistic insights into the enlarged mt-nucleoids observed in HSV-1–infected cells. We discovered that HSV-1 infection degrades TFAM protein in a UL12.5-dependent manner, which could hinder mtDNA replication and nucleoid organization. Exogenously expressed TFAM was degraded as efficiently as the endogenous TFAM in HSV-1–infected cells, whereas *TFAM* transcript levels remained unaltered during HSV-1 infection. TFAM down-modulation was also observed when UL12.5 was expressed outside the context of infection. In both cases, TFAM was partially rescued when bafilomycin A1 was added to the cultures before UL12.5 expression. We also discovered that a second, UL12.5-independent mechanism, down-modulates TFAM in HSV-1 infections when mtDNA is absent, such as in a ρ0 cell line. It is unclear yet if the two mechanisms are active concurrently, where the UL12.5-dependent loss of TFAM acts as the dominant mechanism. It is also possible that HSV-1, under the stress environment of p0 cells, activates another mechanism that causes TFAM down-modulation. These findings highlight the mitochondrial versatility and indicate that different mechanisms likely down-modulate TFAM under different stress conditions. Along with TFAM, mtDNA was also down-modulated in a UL12.5-dependent mechanism, and for this reason UL12.5 was previously used to develop a ρ0 cell line (25). However, UL12.5 expression alone did not cause IFN-β or ISG expression. Instead, we found that the UL12.5-expressing cells mount stronger type I IFN responses in the presence of an innate immunity stimulus such as 2’3’-cGAMP. These findings indicate that UL12.5 likely primes the cells to activate a stress expression profile by disrupting mitochondrial biogenesis. In human cells where HSV-1 productively replicates, like human primary epidermal keratinocytes, fibroblasts, and retinal epithelial cells, this priming does not automatically result in type I IFN gene expression. This is because HSV-1 has evolved elegant mechanisms to evade antiviral responses, and these priming signaling events are likely harnessed to support the infection. However, in non-natural hosts and immune cells UL12.5 likely drives immunity activation. Along with TFAM we also noticed down-modulation of PGC-1α, another critical regulator of mitochondrial biogenesis. This appears to be an essential step in the virus life cycle, as an upregulation of PGC-1α negatively impacts the infection. Several mitochondrial transcription regulators upstream of PGC-1α, like SIRT1 and YY1, are subjugated by HSV-1 for viral gene expression. How changes in these factors influence mitochondria and whether they initiate some of the mitochondrial changes documented during infection is currently unknown.

We also documented UL12.5–independent effects on mitochondria during HSV-1 infection. These effects were observed irrespective of TFAM and mtDNA loss, so they likely precede this loss. Virus entry to the cells was not sufficient to cause these UL12.5-independent effects, indicating that viral gene expression is required. We further discovered changes to mitochondrial membrane proteins that abrogate mitochondrial fusion. The IMM fusion protein OPA1 is converted to shorter forms (S-OPA1) that supports nucleoid structures and the respirasome, but cannot support IMM fusion in the absence of the membrane-bound, long forms (L-OPA1) (16;20;50). This conversion of L- to S-OPA1 was due to OMA1 activation in HSV-1 infected cells. The increased activity of OMA1 was evidenced by its degradation. YME1L1 negatively regulates OMA1 activity in HSV-1–infected cells, since in the absence of YME1L1, the conversion of L- to S-OPA1 was more pronounced. The OMM protein MFN2 is down-modulated in HSV-1-infected cells, which likely disrupts OMM fusion. Mitochondrial footprint and branch length also decrease during infection, a phenotype consistent with mitochondrial fusion disruption. We did not, however, notice any changes in mitochondrial fission proteins in HSV-1-infected cells, indicating that this machinery likely remains functional to produce smaller mitochondria (Figure 10). In support of this, we found that exogenous expression of the long OPA1 isoform 1, which functions to sustain fusion has a negative effect on the infection. Similarly, depletion of OMA1, which is required for the inhibition of fusion negatively impacts the infection, whereas the depletion of YME1L1, a negative regulator of OMA1, promotes the infection. Additionally, we showed the inhibition of fission using a Drp1 inhibitor negatively impacted the infection. Altogether, our data indicate that these mitochondrial membrane protein changes prevent the formation of large (fused) mitochondria, and an extended network, but do not interfere with the formation of smaller mitochondria (Figure 10). The virus could promote these changes because: i) smaller mitochondria can be moved easier to perinuclear sites to supply energy for virus replication and virion assembly; and ii) fusion is an energy demanding process. While our data indicate that the mitochondrial changes benefit the infection, infection of the p0 cell line did not have any impact on HSV-1 infection. The p0 cell line is cultured in the presence of pyruvate and uridine that rescue the metabolic profile of OXPHOS dysfunction. These cells also undergo transcriptional and epigenetic drift to adapt to metabolic changes. Together these changes may mask a potential effect of mtDNA elimination on HSV-1 infection.

**Figure 10:**
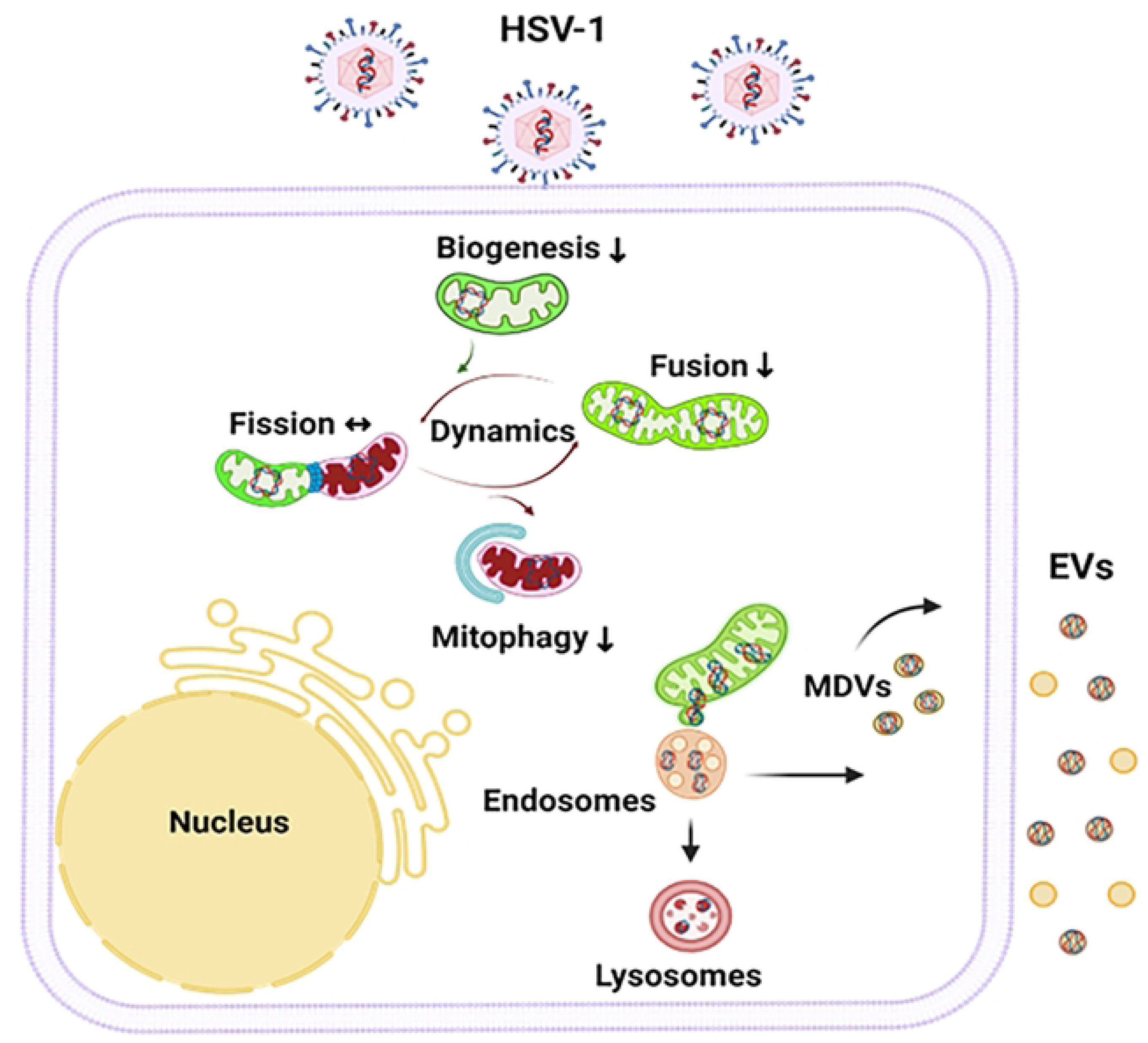
Model summarizing the impact of HSV-1 infection on mitochondria. HSV-1 infection subverts the mitochondrial network to support the infection. Mitochondrial biogenesis and fusion are inhibited during HSV-1 infection. Conversely, mitochondrial fission is preserved, while mitophagy is inhibited to protect mitochondria. Damaged mitochondria content is exocytosed through EVs. These modified mitochondria cluster to perinuclear sites likely supporting virus replication and envelopment.

Several membrane transport complexes are also disrupted in HSV-1 infected cells, particularly in the IMM. The TIM23 protein of the homonymous IMM translocase complex is down-modulated during HSV-1 infection. This effect is likely independent of the changes on the OMM and IMM fusion proteins. TIM23 is critical for importing proteins into mitochondria, including subunits of respiratory enzymes, and its down-modulation is expected to negatively impact mitochondrial protein import (34;35). However, this effect may not be detrimental for mitochondria because most respiratory enzymes have a very long half-life (61;82). We found that exogenous expression of TIM23 had a negative effect on the infection, indicating that the virus likely promotes TIM23 down-modulation in order to support the infection. The ANT2 transporter was also down-modulated late during HSV-1 infection. The ANT2 complex primarily facilitates the transport of ATP from mitochondria to the cytoplasm and the import of ADP to the mitochondrial matrix to be used for ATP production (57). ANT2 also has a role in exporting mitochondrial RNAs to the cytoplasm, where they can activate RNA-sensing pathways (60). It is thus possible that the virus down-modulates ANT2 in order to evade these antiviral responses.

The extensive mitochondrial alterations in HSV-1-infected cells prompted us to investigate if these mitochondria are subjected to mitophagy. This is a mechanism for the removal of irreversibly damaged mitochondria. We determined that such a mechanism is unlikely to eliminate the enlarged mt-nucleoids and other “damaged” mitochondrial content during HSV-1 infection because: i) the mitophagy adaptors are down-modulated before mtDNA/TFAM loss; and ii) mitochondria are used in HSV-1 infected cells. Less invasive mechanisms for selective removal of damaged mitochondrial content exist, involving either a transient contact of mitochondria with endosomes/lysosomes, or a release of mitochondrial-derived vesicles (MDVs) (27;40;41;83). These mechanisms could facilitate the extrusion of mtDNA and other “damaged” mitochondrial content that are then trafficked to lysosomes for degradation, independent of mitophagy. Alternatively, the resulting vesicles could be directed for exocytosis through an EV biogenesis pathway (Figure 10). In human cells, where HSV-1 productively replicates, the autophagy/mitophagy pathways are inhibited by the virus, and lysosomal activity is reduced (39;84-87). Collectively, these findings could explain the increased exocytosis observed in HSV-1 infected cells (73-77;88;89). We also found that both lysosomal degradation and exocytosis have a contributing role in mtDNA/TFAM down-modulation in HSV-1-infected cells. We observed that endosomes intercalate with the mitochondrial network both in infected and uninfected cells, indicating that this represents a canonical pathway for removal of mitochondrial content. However, the proximity of endosomes to mitochondria was more pronounced in infected cells, occasionally resulting in transient interactions between the two organelles, aka “kissing events”. Despite these findings, a role for mitochondrial nucleases and proteases in mtDNA/TFAM degradation cannot be excluded.

While HSV-1 infection impacts mitochondrial dynamics, mitochondria retain certain functions. Basal respiration of the cells is not altered up to late hours post-infection, and there is an increase in spare respiratory capacity. This is likely an adaptation of the infected cells to their increased energy demand. We also noticed an increase in basal ECAR and maximal glycolysis, an indication that the virus needs rapid energy production, and nucleotide synthesis to support its replication. The mitochondrial membrane potential Δψm was preserved in HSV-1-infected cells. This is critical as the disruption of Δψm inhibits viral gene expression, decreases progeny virus production, and promotes mitochondrial fission and fragmentation. Finally, we determined that mitochondrial superoxide production remains unaltered between infected and uninfected cells.

Disruption of mitochondrial homeostasis is a hallmark of almost all neurodegenerative diseases. Because of their high energy requirements, neurons are particularly vulnerable to injury and death from dysfunctional mitochondria. Because they are post-mitotic, neurons also accumulate mitochondrial damage with age. HSV-1 infection has been linked to neuronal dysfunction, in part due to repeated “clinically silent” reactivations in the brain that may promote neurodegenerative processes (90–92). Defining the mechanisms that disrupt mitochondrial homeostasis during HSV-1 infection can yield insight into the molecular basis of virus pathogenesis.

## Acknowledgements

We would like to thank Dr. Sandra Weller (University of Connecticut) for kindly sharing the AN-1, ANF-1, KOS 37 strains, the Vero 6-5 cell line, and the UL12/UL12.5 antibody. Also, we would like to thank Dr. James R. Smiley (University of Alberta) for kindly sharing the KOS37 UL98-SPA virus. Further, we would like to thank the KU Alzheimer’s Disease Research Center (P30AG072973) Biomarker Core assisted the Seahorse analysis, and the KUMC Integrative Imaging Core for facilitated the microscopy studies. This study was supported by the NIAID R01AI162784. The KUMC Leica TCS SP8 STED is supported through the NIH S10 OD 023625 and the Nikon CSU-W1 SoRa through the NIH S10 OD032207.

## Supplemental information

**Figure S1: ΔUL47 virus cannot evade antiviral responses. A.** hTERT-HEL cells were infected with HSV-1(F) or ΔUL47 virus (0.01 PFU/cell). The cells were harvested at 3, 24, and 48 hpi and progeny virus production was determined by plaque assays in Vero cells. **B.** hTERT-HEL cells were infected with HSV-1(F) or ΔUL47 virus (0.1 PFU/cell) or left uninfected. The cells were harvested at 14 hpi, total RNA was extracted and type I IFN gene expression was quantified by RT-qPCR. 18S rRNA was used for normalization. **C.** hTERT-HEL cells and ATG5 KD derivatives were infected with HSV-1(F), ΔUL47 virus (2.5 PFU/cell) or left uninfected. The cells were harvested at 3 hpi and equal amounts of proteins from total cell lysates were analyzed for LC3 and β-actin. Increased levels of LC3-II over LC3-I is an indication of autophagy activation. Autophagy is not observed in ATG5 KD cells.

**Figure S2: Effect of HSV-1 infection on mitochondrial transcription factors. A.** hTERT-HEL cells were infected with HSV-1(F) (2.5 PFU/cell) or left uninfected. The cells were harvested at 3, 6, and 14 hpi. Total RNA was extracted using Trizol, converted to cDNA, and quantified by qPCR using TFAM and PGC-1α specific probes. **B.** hTERT-HEL cells were infected with HSV-1(F) and ΔUL47 (2.5 PFU/cell), the cells were harvested at 3, 14, and 24 hpi and equal amounts of proteins from total cell lysates were analyzed PGC-1α, TFAM, NRF1, and SIRT1. ICP0 served as a control for the infection, and β-actin as a loading control. Quantification of PGC-1α protein signal from three independent experiments was done using ImageJ. **C.** hTERT-HEL cells were treated for 2 h with different doses of ZLN005 followed by HSV-1(F) infection (0.5 PFU/cell). The cells were harvested at 12 hpi and equal amounts of proteins from total cell lysates were analyzed for ICP0, ICP4, ICP8, gD, VP16, and β-actin that served as loading control. **D.** An hTERT-HEL SIRT1 KD cell line was developed with the aid of a lentiviral vector carrying a specific shRNA against SITR1. hTERT-HEL cells and SIRT1 KD derivatives were infected HSV-1(F) (0.01 PFU/cell). The cells were harvested at 3, 24, and 48 hpi and progeny virus production was quantified by plaque assays. **E.** hTERT-HEL YY1 KD cell lines were developed with the aid of lentiviral vectors carrying specific shRNAs against YY1. hTERT-HEL cells and two YY1 KD derived clones were infected with HSV-1(F) (2.5 PFU/cell). The cells were harvested at 16 hpi and equal amounts of proteins from total cell lysates were analyzed for TFAM and YY1. ICP0 and VP16 were used as controls for the infection, and β-actin as a loading control. **F.** hTERT-HEL cells and YY1 KD derivatives were infected HSV-1(F) (0.01 PFU/cell). The cells were harvested at 3, 24, and 48 hpi and progeny virus production was quantified by plaque assays. All statistical analyses were performed with two-tailed Student’s *t*-test from three biological replicates. P<0.05*, P<0.01**, P<0.001***, ns= not significant.

**Figure S3: mtDNA is not required for viral gene expression. A.** HaCaT cells were infected with KOS37, KOS UL98, AN-1, ANF-1, and HSV-1(F) (5 PFU/cell) or left uninfected. The cells were harvested at 20 hpi and equal amounts of proteins from total cell lysates were analyzed for TFAM, OPA1, and TIM23. VP16 was used as controls for the infection and β-actin as a loading control. Quantification of mtDNA was done as described earlier. **B.** hTERT-HEL cells were infected with HSV-1(F) (5 PFU/cell) or left uninfected. Cells were treated with different doses of the LonP1 inhibitor that was added to the cultures at 3 h after infection. The samples were harvested at 14 h post-infection and equal amounts of proteins from total cell lysates were analyzed for TFAM, OMA1, NBR1, p62/SQSTM1, and several viral proteins. β-actin served as a loading control. **C.** SH-S55Y cells and the p0 derivatives were infected with HSV-1(F) (0.5 PFU/cell) or left uninfected. The cells were harvested at 6, 9, and 12 h post-infection and equal amounts of proteins from total cell lysates were analyzed for ICP4 and VP16. β-actin served as a loading control.

**Figure S4: HSV-1 UL12/UL12.5 deficient viruses do not activate type I IFN responses in human cells where HSV-1 productively replicates. A.** hTERT-HEL cells were infected with KOS37, KOS UL98, AN-1, ANF-1, and HSV-1(F) (0.1 PFU/cell). At 2 h post-infection 2’3’-cGAMP (12 μM) was added to the cultures. Samples were harvested at 14 hpi, total RNA was extracted, and quantification of *ISG56*, *ISG15*, *ISG20*, *IFIH1*, and *IL-6* transcripts was done by RT-qPCR. Normalization was done against 18S rRNA. Experiments were performed three independent times and analyzed for statistical significance with two-tailed Student’s *t*-test. P<0.05*, P<0.01**, P<0.001***, P<0.0001****, ns= not significant. **B.** hTERT-HEL cells were infected with KOS37, KOS UL98, AN-1, ANF-1, and HSV-1(F) (5 PFU/cell). Samples were harvested at 14 hpi and equal amounts of proteins from total cell lysates were analyzed for LC3 and OPTN. ICP0 was used as a control for the infection and β-actin as a loading control. The ratio of LC3-II (lipidated) to LC3-I (non-lipidated) is depicted. **C.** hTERT-HEL cells were infected with HSV-1(F) (5 PFU/cell). BafA1 (15 μM) and MG132 (20 μM) were added to the cultures at 4 h and 6 hpi. Samples were harvested at 14 hpi and equal amounts of proteins from total cell lysates were analyzed for TFAM, TIM23, OMA1, NBR1, and MFN2. ICP4 and VP16 were used as controls for the infection and β-actin as a loading control.

**Figure S5: Mitochondria and endosomes display increased proximity during HSV-1 infection. A.** ARPE-19 cells were co-transfected with a Twinkle and RFP-Rab7, or a LAMP1-RFP–expressing plasmid. At 16 h post-transfection, the cells were infected with HSV-1(F) (10 PFU/cell) or left uninfected. The cells were fixed at 16 hpi and double-stained with a V5 antibody to detect Twinkle and an ICP4 antibody to visualize the infection. Images were acquired with a Leica TCS SP8 confocal microscope using the LasX software provided by Leica. **B.** ARPE-19 cells were co-transfected with a Twinkle and mCherry-Rab5–expressing plasmid and infected with HSV-1(F) as above. The samples were fixed at 16 hpi and double-stained with a V5 antibody to visualize Twinkle, and an ICP4 antibody to visualize the infection. Images were acquired with a Leica TCS SP8 confocal microscope and analyzed with the LasX software. **C.** Transfections in ARPE-19 cells were done as in panel A, with a TOM20-EGFP and either a mCherry-Rab5 or an RFP-Rab7–expressing plasmid. At 16 h post-transfection, the cells were infected with HSV-1(F) (10 PFU/cell) or left uninfected. The samples were fixed at 16 hpi, stained with an ICP4 antibody to visualize the infection, and imaged, as in panel A.

## Materials and Methods

### Cell lines and viruses

hTERT-HEL cells (immortalized human embryonic lung fibroblasts, human telomerase reverse transcriptase [hTERT] transformed) were cultured in Dulbecco’s modified Eagle’s medium supplemented with 10% fetal bovine serum (FBS). ARPE-19, HEp-2, Vero, U2OS and 293T cells (ATCC) were cultured according to the manufacturer’s instructions. HSV-1(F) is a limited passage isolate, as described before (93). Properties of ΔICP0 (R8501) and ΔUL47 viruses were described before (42;94-96). The UL12/UL12.5-null virus (AN-1) that does not express UL12 or UL12.5, and the UL12 mutant virus (ANF-1) that produces only UL12.5, were a gift from Dr. Weller (63;97). The KOS strain of HSV-1 is the parental virus for AN-1 and ANF-1. The Vero-derived cell line 6-5 was used to complement UL12 mutant viruses and was propagated in the medium supplemented with 250 ug/ml of G418 (gift from Dr. Weller, University of Connecticut) (63;97). The KOS37 UL98-SPA is a UL12/UL12.5-null that expresses a C-terminally SPA-tagged version of the UL12 ortholog HCMV UL98 (gift from Dr. Smiley, University of Alberta)(66). The SH-SY5Y (p+) and the SH-SY5Y (ρ0) cell lines were provided by Dr. Swerdlow (University of Kansas Medical Center) (68;69). The SH-SY5Y ρ0 cell line was maintained in DMEM supplemented with 10% FBS (filtered sterile), pyruvate (1 mM) and uridine (0.25 mM).

### Plasmids

The pLenti-CMV-Puro-TFAM plasmid was developed by cloning the TFAM ORF into the BamHI-SalI sites of the pLenti-CMV-GFP vector. The TFAM ORF was obtained from the pcDNA3 TFAM-mScarlet plasmid by PCR (forward: 5’ CC AAG CTT GAT ATC ATG GCG TTT CTC CGA 3’ and reverse: 5’ GG AAG CTT GAT ATC TTA ACA CTC CTC AGC ACC 3’) followed by digestion with EcoRV. The pLenti-CMV-Puro-3XHA-EGFP-OMP25 plasmid was obtained by cloning the EcoRI-NotI fragment from pMXs-3XHA-EGFP-OMP25 into the BamHI-SalI sites of the pLenti CMV GFP vector. The pcDNA3.1-Zeo(+)-UL12.5-EGFP plasmid was developed by cloning the UL12.5-EGFP ORF into the EcoRV site of the pcDNA3.1-Zeo(+) plasmid. The pcDNA3.1-Zeo(+)-UL12.5-Flag plasmid was developed by cloning the UL12.5-Flag ORF into the EcoRV site of pcDNA3.1-Zeo(+) plasmid. The pLenti-mCherry-Mango II x 24-UL12.5-Flag and UL47-Flag plasmids were developed by cloning the UL12.5-Flag ORF and the UL47-Flag ORF into the BamHI-NheI sites of pLenti-mCherry-Mango II x 24. The UL12.5-EGFP and UL12.5-Flag ORFs were custom synthesized through GenScript. All plasmids were verified by sequencing.

### Development of stable cell lines and transfection assays

Plasmids carrying shRNAs for the depletion of human ATG5, SIRT1, and YY1 were purchased from Sigma. For overexpressing and inducible cell lines we developed several plasmids described earlier. To produce lentiviruses, HEK-293T cells seeded in a F25 cm^2^ flask at a 60% confluency were cotransfected with the plasmid carrying the shRNA, the Gag-Pol–expressing plasmid (pCMV-dR8.2 dvpr), and the VSV-G–expressing plasmid (pCMV-VSV-G) using Lipofectamine 3000 (5 μg total, ratio 7:7:1), according to manufacturer’s instruction (Thermo Fisher Scientific). At 48 h after transfection, the supernatant from the cultures was collected, filtered through 0.45-μm-pore-size filters, and used to transduce hTERT-HEL cells in the presence of polybrene (10 μg/ml). Puromycin selection (2 μg/mL) was initiated 24 h after exposure to lentiviruses and continued until only resistant clones emerged. The clones of hTERT-HEL cells, with the greater depletion in the protein of interest, were used. The stable lines were maintained in the presence of puromycin, which was removed 24 h before assays were performed.

For other transfections, the ratio of plasmid(s) to Lipofectamine 3000 for a given number of cells was calculated according to the manufacturer’s instructions. All plasmids used in this study are listed in the key resources table. All siRNAs and control siRNAs (scrambled) were transfected at a dose of 200 ng/ 10^6^ cells for 72 hours prior to infection.

### Extracellular vesicle purification

Isolation of EVs from infected or uninfected cells was done as previously described, with minor modifications (73;75;77). Briefly, hTERT-HEL cells were infected with HSV-1(F) (0.5 PFU/cell) or remained uninfected. The supernatant was collected at 24 h post-infection, centrifuged at 1,000x *g* for 10 min and at 2,000x *g* for 20 min to remove floating cells and larger cell debris. The supernatant was filtered through a 0.45-μm-pore-size filter and concentrated through centricon devices (molecular mass cutoff, 100 kDa; Centricon Plus 70), according to the manufacturer’s instructions (Millipore). The samples were then subjected to 15,000x *g* centrifugation for 2 h to remove virions and heavier particles. Total EVs were obtained by concentrating the supernatant either through ultracentrifugation (100,000x *g*, 3 h) to analyze for proteins, or through 100 kDa filters (Vivaspin20) to analyze for DNA. These EVs were loaded on top of an iodixanol-sucrose gradient that ranged from 6% to 18%, in 1.2% increments. The 60% iodixanol was diluted in 10 mM Tris, pH 8, and 0.25 M sucrose. Samples were centrifuged using an SW41Ti rotor for 2 h at *r_max_*250,000 ×*g* at 4°C in a Beckman Coulter Optima XPN-80 ultracentrifuge. Fractions (2 ml) were collected from the top to the bottom of the gradient for further analysis.

### Western blot analysis

Cells were solubilized in triple detergent buffer (50 mM Tris-HCl [pH 8], 150 mM NaCl, 0.1% sodium dodecyl sulfate, 1% Nonidet P-40, 0.5% sodium deoxycholate, 100 mg/ml of phenylmethylsulfonyl fluoride) supplemented with phosphatase inhibitors (10 mM NaF, 10 mM β-glycerophosphate, 0.1 mM sodium vanadate) and protease inhibitor cocktail (Sigma), then briefly sonicated. Protein concentration was determined with the Bradford method (Bio-Rad Laboratories). The antibodies used and their dilutions are listed in the key resources table. Proteins were visualized with SuperSignal West Pico PLUS Chemiluminescent Substrate (Thermo Fisher Scientific) using the ImageQuant 800 Western blot imaging system (Amersham).

### Immunofluorescence analysis

The procedures were described elsewhere (74). Briefly, the cells were fixed in 4% paraformaldehyde (PFA), permeabilized, blocked with PBS–TBH solution (consisting of 0.1% Triton X-100 in PBS, 10% horse serum, and 1% BSA), and incubated with primary antibodies (key resources table) diluted in PBS–TBH. The cultures were washed 3x with PBS–TBH and incubated with appropriate Alexa-Fluor-conjugated secondary antibodies, diluted in PBS–TBH. After several rinses, first with PBS–TBH and then with PBS, the samples were mounted in mounting medium (plain or with DAPI) and examined with a Leica TCS SP8 STED super resolution microscope. Images were acquired at 100x magnification, pinhole at 1, a resolution of 2048×2048 pixels for 2D images, and 1024×1024 pixels for Z-stacking. For 3D images, approximately 20-30 images were acquired with a z-step set at 0.30 μM. The LasX software was used for image analysis and for orthogonal section view.

### Viral genome, mtDNA, and mRNA quantification

Total DNA was extracted from the cells using the NucleoSpin DNA RapidLyse kit (Macherey-Nagel). mtDNA was quantified with a human mitochondrial DNA (mtDNA) monitoring primer set (Takara), which quantifies the relative number of copies of human mtDNA using nuclear DNA (nDNA) content as a standard. To quantify HSV-1 genome, we used primer pairs targeting the area of the viral genome encoding gI (see table of key resources). For normalization, we used primers targeting β-actin DNA. Both primer pairs have been described before (29, 30).

Quantification of transcripts was performed in a QuantStudio5 Real-Time PCR system (Applied Biosystems), using PowerUp^TM^ SYBR^TM^ Green Master Mix or TaqMan^TM^ Universal Master mix with UNG (Applied Biosystems) according to manufacturer’s instructions. Pre-designed probes (FAM™/MGB) for the human TFAM and PGC-1α transcripts were obtained through Thermo Fisher Scientific. The 18S rRNA primers (Universal Primers, Ambion) were used for normalization. Other primer pairs are listed under the key resources table.

### Seahorse Mito Stress test

hTERT-HEL cells (15,000) were seeded in a Seahorse cell culture plate (Agilent) and cultured in 10% fetal bovine serum supplemented Dulbecco’s modified Eagle’s Medium. Mitochondria stress test was performed using a Seahorse XF Pro Analyzer (Agilent), and the cell oxygen consumption rate (OCR) and extracellular acidification rates (ECAR) were quantified. The assay was performed in XF DMEM Base Medium (Agilent) supplemented with 1 mM pyruvate, 1 mM glutamine, and 2.5 mM glucose (Assay media). Different substrates prepared in the assay media were successively injected through the injection port to measure OCR or ECAR. Arrangement of substrate in the injection port were as following: A) 2 µM oligomycin, B) 1 µM FCCP, C) 1 µM antimycin A and 50 nM rotenone, and D) none. Measurements were taken 3 times for 3 min, after 3 min mix post-injection. Eight biological replicates were analyzed using the Agilent Seahorse Wave pro 10.2.1 and GraphPAD Prism 10. Reads were normalized to 1,000 cells. The P values were calculated using a two-tailed Student’s *t*-test. P <0.05*, P<0.01**, P<0.001***.

### Tetramethylrhodamine, Methyl Ester, Perchlorate (TMRM) and MitoSOX Red staining

hTERT-HEL cells seeded in 96-well plates were infected with HSV-1(F) (2.5, 5.0, and 10 PFU/cell). At 8 and 16 h pi cells were incubated in 100 nM of TMRM dye (T668; Thermo Fisher Scientific) for 30 min at 37 °C or 50 μM MitoSOX Red for 45 min at 37°C. The cells were washed 3x with PBS and 100 μl of culture media (DMEM + 10% FBS) was added after the final wash. Live cell images were obtained at 20x magnification (intensity= 10, integration time 31 msec, and camera gain 24) using Cytation 7 (Agilent). Cells were also stained with Hoechst 33342 for cell count. Mean fluorescence intensity (MFI) of the images was measured using the Gen5 3.13 software. Three independent experiments were performed, and two images were taken per condition from each experiment. To calculate the average TMRM MFI, the MFI per cell was calculated by dividing fluorescent intensity of the image with the total number of cells in the image. The average MFI per cell and experiment was calculated and normalized to 100 cells as follows: MFI per 100 cells per experiment from three independent experiments were averaged to finally determine the final TMRM MFI. The P values were calculated using a two-tailed Student’s *t*-test. P < 0.05*, P<0.01**, P<0.001***.

### Analysis of mitochondrial morphology, MitoTtracker Red, and LysoTracker Red staining

hTERT-HEL cells were cultured in 5-well confocal slides (Electron Microscopy Sciences) in DMEM 10% FBS. Cells were infected with HSV-1(F) (5 PFU/cell), and at 16 h post-infection the cells were treated with MitoTracker Red (100 nM) for 30 min at 37°C. Then, the cells were washed 3x with PBS to remove excess dye, fixed in 4% PFA, and stained with ICP4 antibody to visualize the infection. Images were acquired using a Leica TCS SP8 STED confocal microscope at 100X magnification (15.6% laser intensity). Mean fluorescence intensity (MFI) and total area per cell were measured using ImageJ. Total fluorescence intensity (TFI) of a cell was calculated by multiplying the MFI of the cell with the total area of the cell. The average TFI from uninfected (n=12) and HSV-1(F)–infected cells (n=30) was plotted in a bar graph. For mitochondrial morphology analysis, we used the FiJi distribution of ImageJ with the Macro tool MiNA (Mitochondrial Network Analysis), as detailed before (48). The P values were calculated using a two-tailed Student’s *t*-test. P < 0.05*, P<0.01**, P<0.001***. LysoTracker Red (100 nM) was added to live cells for 1 h at 37 ^0^C, and excess dye was removed with PBS prior fixation.

### Key resources table

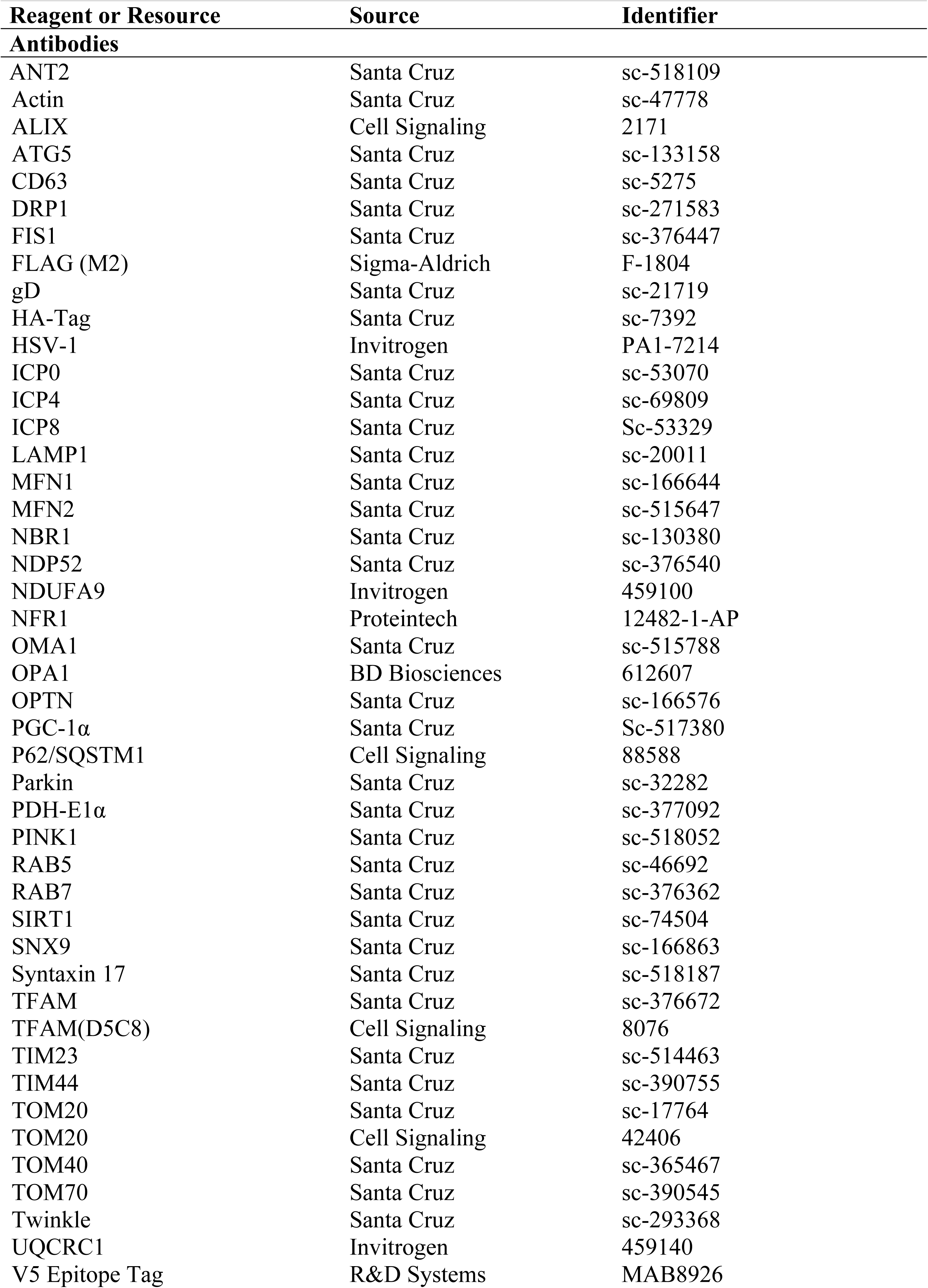
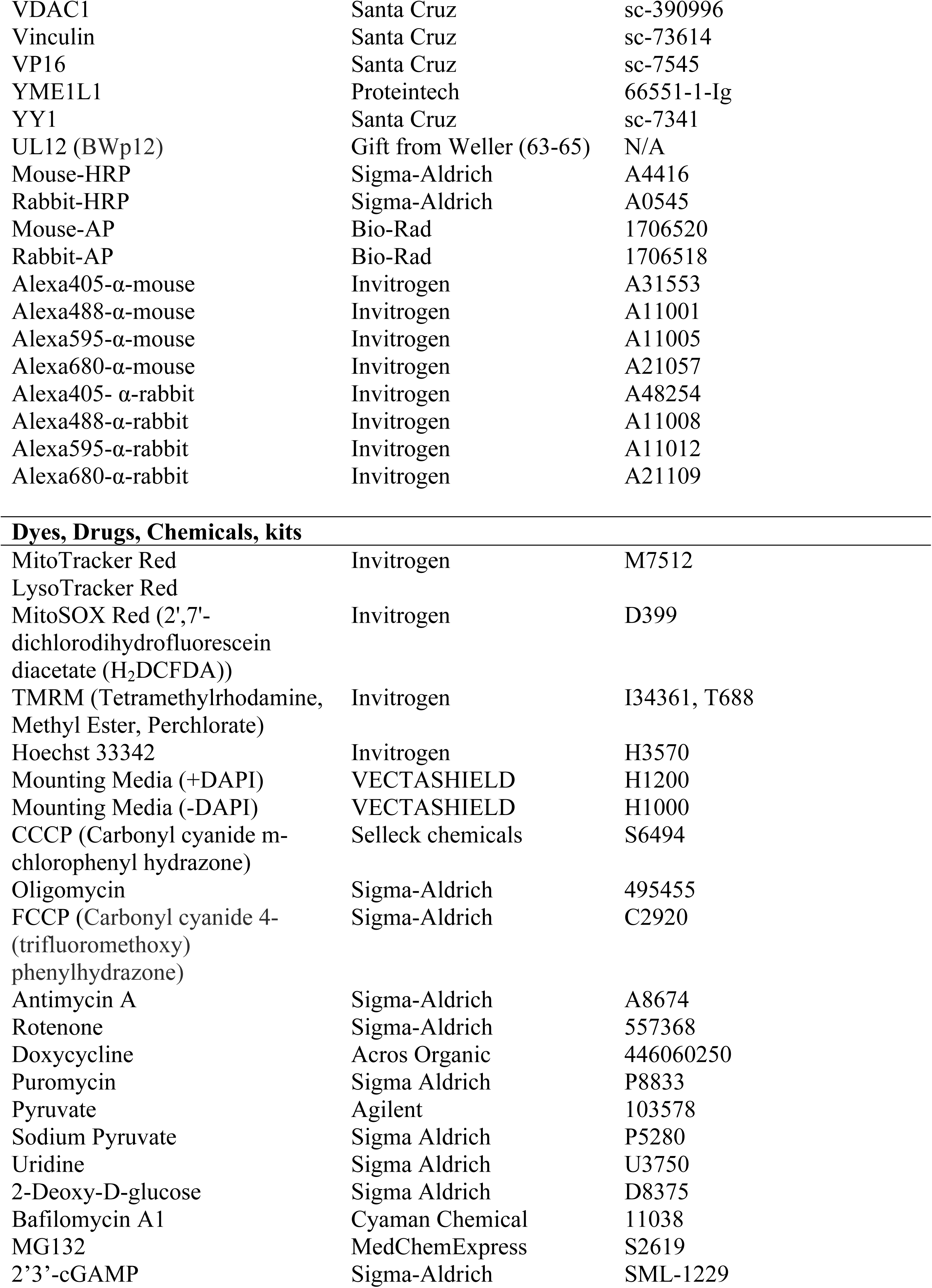
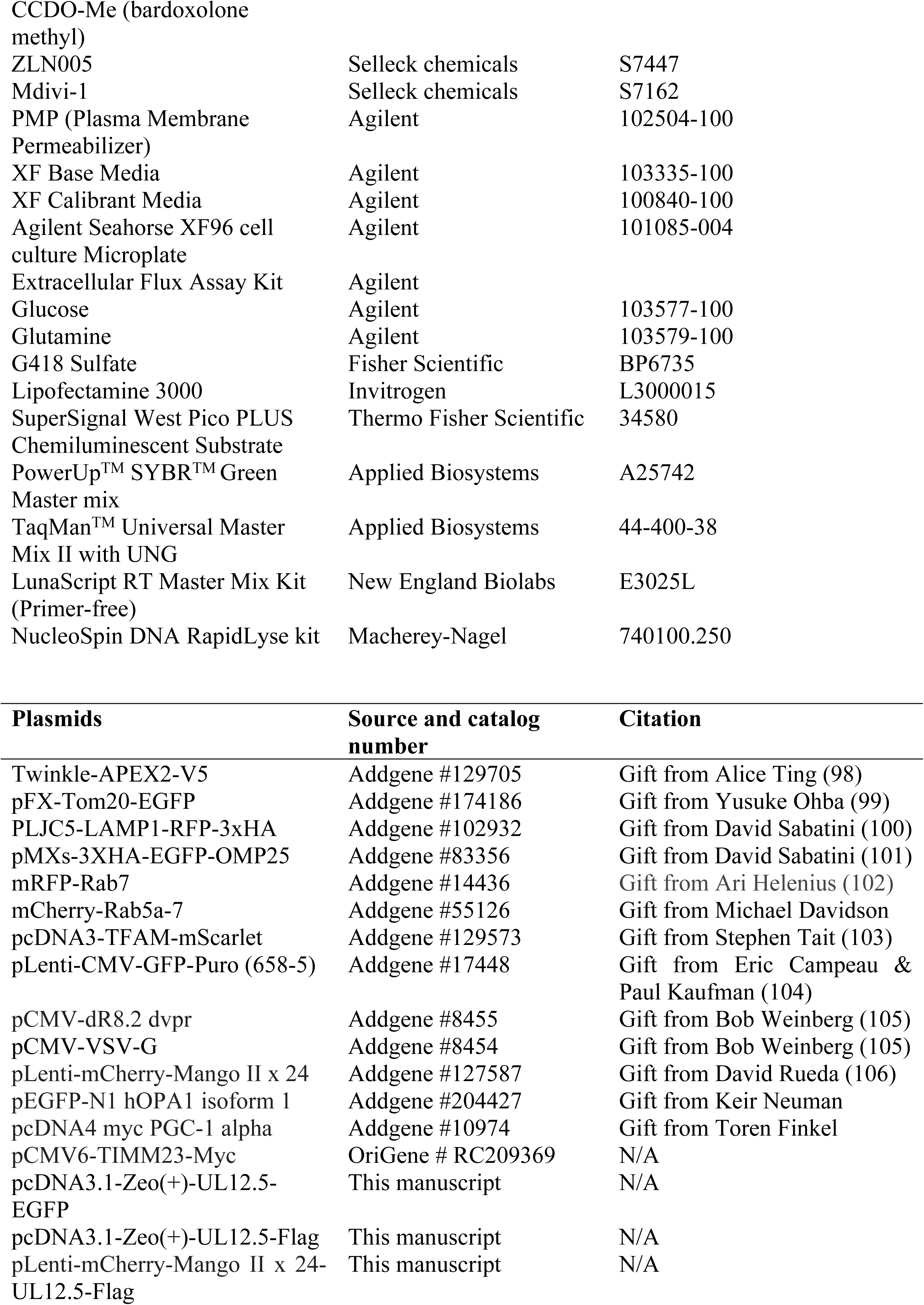
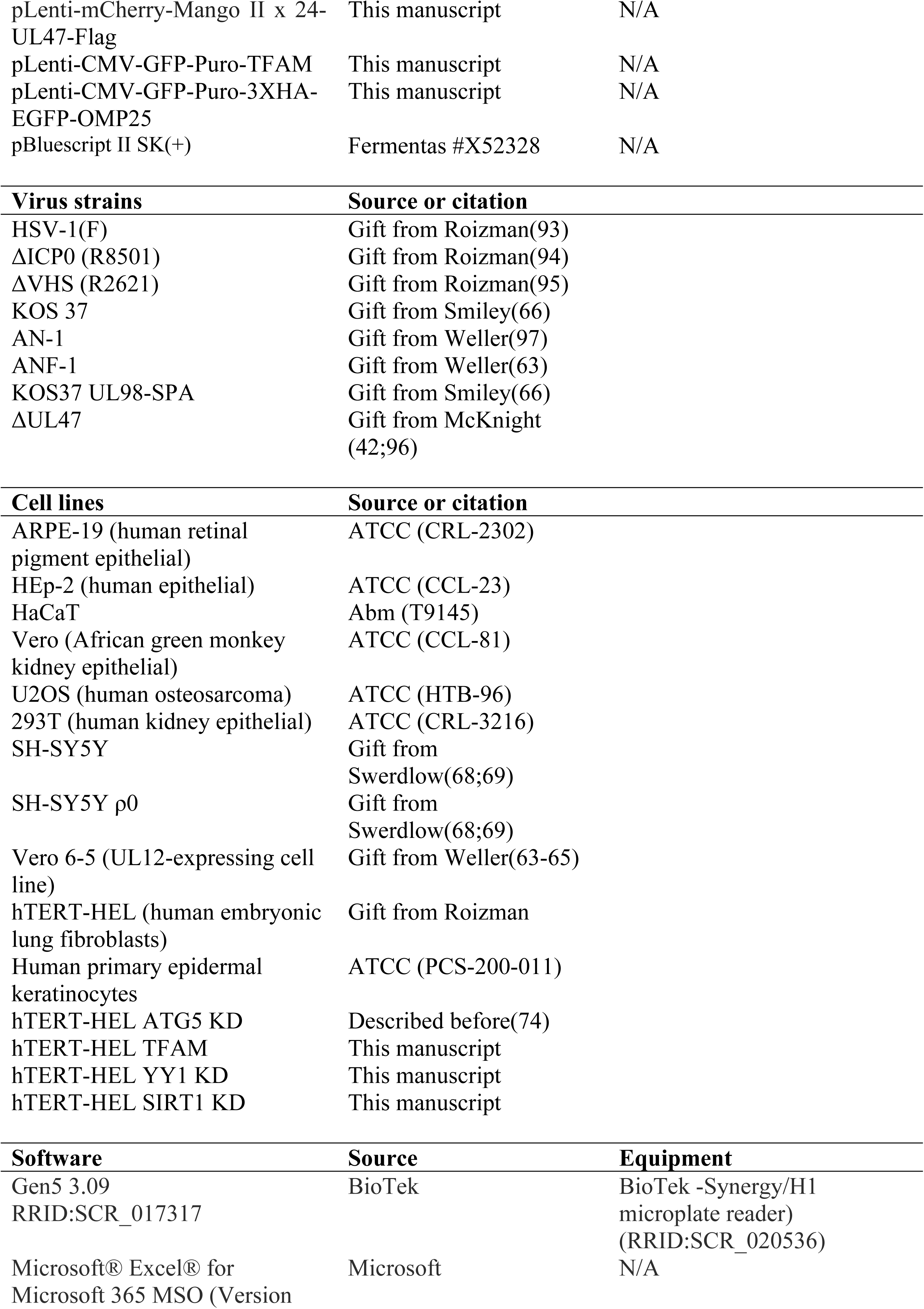
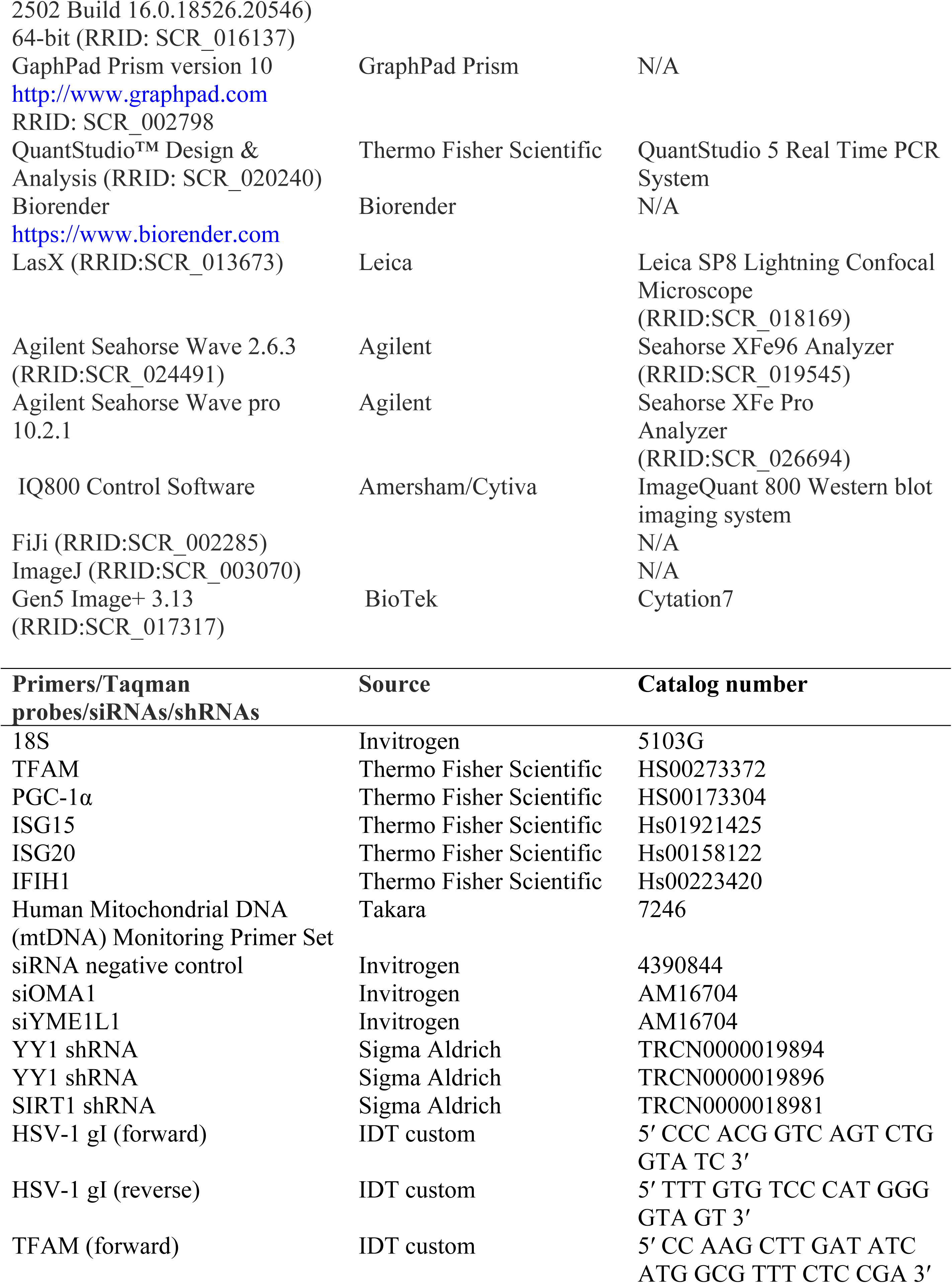
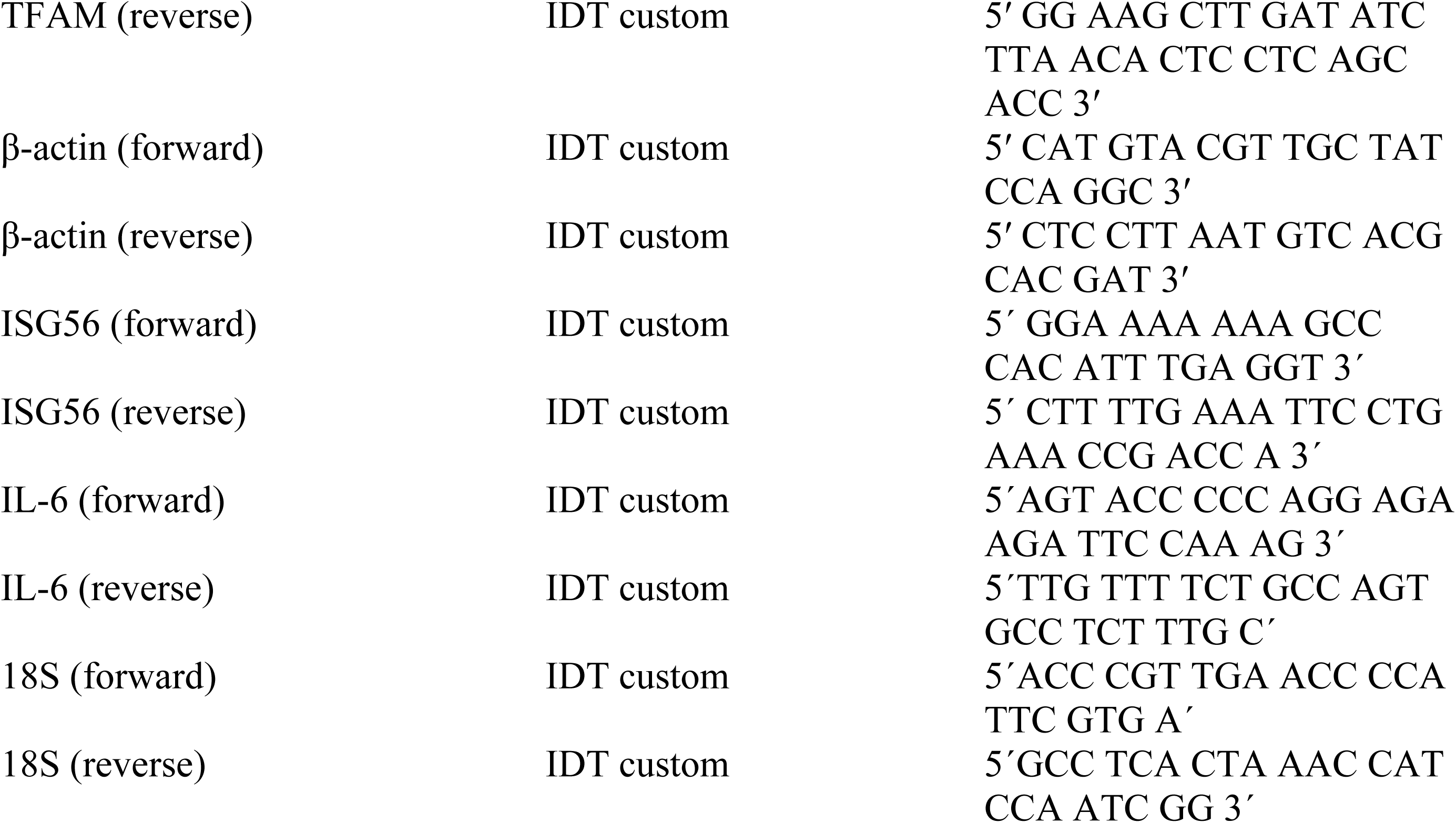

## Notes

### Competing Interest Statement

The authors have declared no competing interest.

